# Mitohormesis reprograms macrophage metabolism to enforce tolerance

**DOI:** 10.1101/2020.10.20.347443

**Authors:** Greg A. Timblin, Kevin M. Tharp, Breanna Ford, Janet M. Winchenster, Jerome Wang, Stella Zhu, Rida I. Khan, Shannon K. Louie, Anthony T. Iavarone, Johanna ten Hoeve, Daniel K. Nomura, Andreas Stahl, Kaoru Saijo

**Affiliations:** Department of Molecular and Cell Biology, University of California, Berkeley, Berkeley, CA 94720; Center for Bioengineering and Tissue Regeneration, Department of Surgery, University of California San Francisco, San Francisco, CA 94143; Department of Nutritional Sciences and Toxicology, University of California, Berkeley, Berkeley, CA 94720; Novartis-Berkeley Center for Proteomics and Chemistry Technologies and Department of Chemistry, University of California, Berkeley, Berkeley, CA 94720; California Institute for Quantitative Biosciences (QB3), University of California, Berkeley, Berkeley, CA 94720; QB3/Chemistry Mass Spectrometry Facility, University of California, Berkeley, Berkeley, CA 94720; Department of Molecular and Medical Pharmacology, Crump Institute for Molecular Imaging and UCLA Metabolomics Center, University of California, Los Angeles, Los Angeles, CA 90095; Helen Wills Neuroscience Institute, University of California, Berkeley, Berkeley, CA 94720

## Abstract

Macrophages generate mitochondrial reactive oxygen and electrophilic species (mtROS, mtRES) as antimicrobials during Toll-like receptor (TLR)-dependent inflammatory responses. Whether mitochondrial stress caused by these molecules impacts macrophage function is unknown. Here we demonstrate that both pharmacologically- and lipopolysaccharide (LPS)-driven mitochondrial stress in macrophages triggers a stress response called mitohormesis. LPS-driven mitohormetic stress adaptations occur as macrophages transition from an LPS-responsive to LPS-tolerant state where stimulus-induced proinflammatory gene transcription is impaired, suggesting tolerance is a product of mitohormesis. Indeed, like LPS, pharmacologically-triggered mitohormesis suppresses mitochondrial oxidative metabolism and acetyl-CoA production needed for histone acetylation and proinflammatory gene transcription, and is sufficient to enforce an LPS-tolerant state. Thus, mtROS and mtRES are TLR-dependent signaling molecules that trigger mitohormesis as a negative feedback mechanism to restrain inflammation via tolerance. Moreover, bypassing TLR signaling and pharmacologically triggering mitohormesis represents a novel anti-inflammatory strategy that co-opts this stress response to impair epigenetic support of proinflammatory gene transcription by mitochondria.

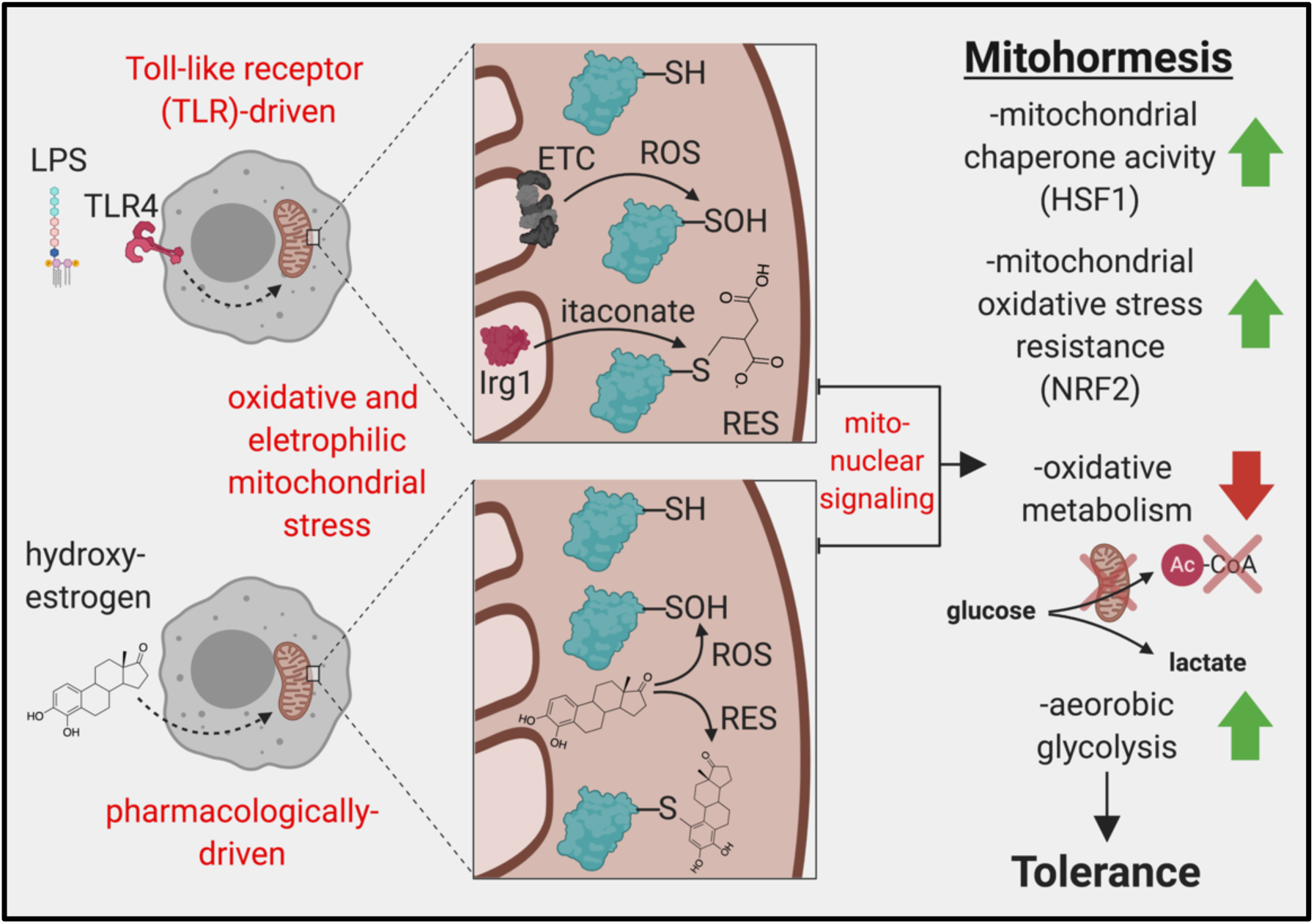

## Introduction

Macrophage inflammatory responses are important for host defense, but if not tightly controlled, can be detrimental to the host in acute and chronic inflammatory diseases^1, 2^. Macrophage tolerance evolved to protect the host from overproduction of inflammatory mediators; however, this immunoparalyzed state impairs the ability of macrophages to clear pathogens, tumors, and perform tissue homeostatic functions^3-6^. Thus, understanding how the balance between responsiveness versus tolerance is regulated has far-reaching implications in health and disease.

Mitochondria are critical for macrophage-mediated immunity^7^. Upon encountering a pathogen, toll-like receptor (TLR) ligands such as lipopolysaccharide (LPS) increase macrophage production of mitochondrial reactive oxygen species (mtROS) for bactericidal purposes^8^. Moreover, TLR-driven expression of *Irg1* results in TCA cycle remodeling and production of itaconate^9^, a mitochondrial reactive electrophilic species (mtRES)^10^ with antimicrobial properties^11^. Rapid production of mtROS and mtRES following TLR engagement coincides with acute oxidative damage^12^, glutathione (GST) depletion^13^, and activation of cytoprotective genes (e.g. *Nfe2l2, Atf4*)^12, 13^, suggesting oxidative and electrophilic mitochondrial stress is caused by local production of these reactive molecules. Whether this acute mitochondrial stress impacts macrophage function long-term is unknown.

Mitochondrial integrity is closely monitored by quality control systems. This includes nuclear-encoded transcriptional factors that detect signs of mitochondrial stress (e.g. increased mtROS^14, 15^, impaired mitochondrial protein import and folding^16^) and alter gene expression via retrograde mito-nuclear signaling. Most thoroughly studied in model organisms such as *Caenorhabditis elegans* and *Saccharomyces cerevisiae*, stress-induced activation of these transcription factors can promote persistent cyto- and mito-protective adaptations and stress resistance in a process known as mitohormesis, which influences organismal metabolism, health, and longevity^17-19^. Whether mitohormesis influences immunity is unknown.

Here we demonstrate that both pharmacological and TLR-driven mitochondrial stress triggers mitohormesis in macrophages. Mitohormetic adaptions, including enhanced mitochondrial proteostasis, mitochondrial oxidative stress resistance, and the suppression of oxidative mitochondrial metabolism, leave macrophages in an immunoparalyzed, LPS-tolerant state. Furthermore, by pharmacologically triggering mitohormesis with reactive, lipophilic small molecules that mimic the actions of endogenously-generated mtROS and mtRES, we show the transition to an LPS-tolerant state can occur independent of TLR signaling and other mechanisms previously proposed to enforce tolerance. Thus, mtROS and mtRES trigger mitohormesis as a negative feedback mechanism to restrain macrophage inflammation via tolerance, and this process can be exploited therapeutically to counteract acute and chronic inflammation.

## Results

### Hydroxyestrogens are anti-inflammatory in macrophages *in vitro*

Biological sex affects immune responses^20^, and estrogens have immunomodulatory effects on macrophages^21, 22^. Recent evidence suggests estrogens can have direct, estrogen receptor (ER)-independent effects on mitochondrial function^23^, which controls macrophage inflammatory responses^7^. While many studies focus on 17β-estradiol (E2), the most abundant estrogen, metabolism can alter the structure and immunomodulatory properties of sterols^24, 25^. We screened a panel of 14 endogenous estrogens (**Supplementary Table 1**) to identify metabolites with anti-inflammatory activity in macrophages that might have therapeutic utility. Bone marrow-derived macrophages (BMDMs) were pretreated with estrogen metabolites for 1 hour, followed by LPS stimulation for 6 hours and *Nos2* quantitative real-time PCR (qPCR). This revealed the hydroxyestrogens 2-hydroxyestrone (2-OHE1), 4-hydroxyestrone (4-OHE1), and 2-hydroxyestradiol (2-OHE2), significantly repressed *Nos2* induction, while other estrogens including E2 and 16-Epiestriol (16-Epi) lacked this activity (**Fig.1A**). RNA-seq identified 253 common genes repressed by individual hydroxyestrogen pretreatment in LPS-stimulated BMDMs (**Supplementary Table 2**). Gene Ontology (GO) analysis of these genes revealed enrichment for categories including “Inflammatory response” and “Cytokine production” (**Fig.1B**). Hierarchical clustering of the RNA-seq data revealed a broad set of proinflammatory cytokines and chemokines repressed by hydroxyestrogens (**Fig.1C)**, which were validated by qPCR

**Figure 1.**
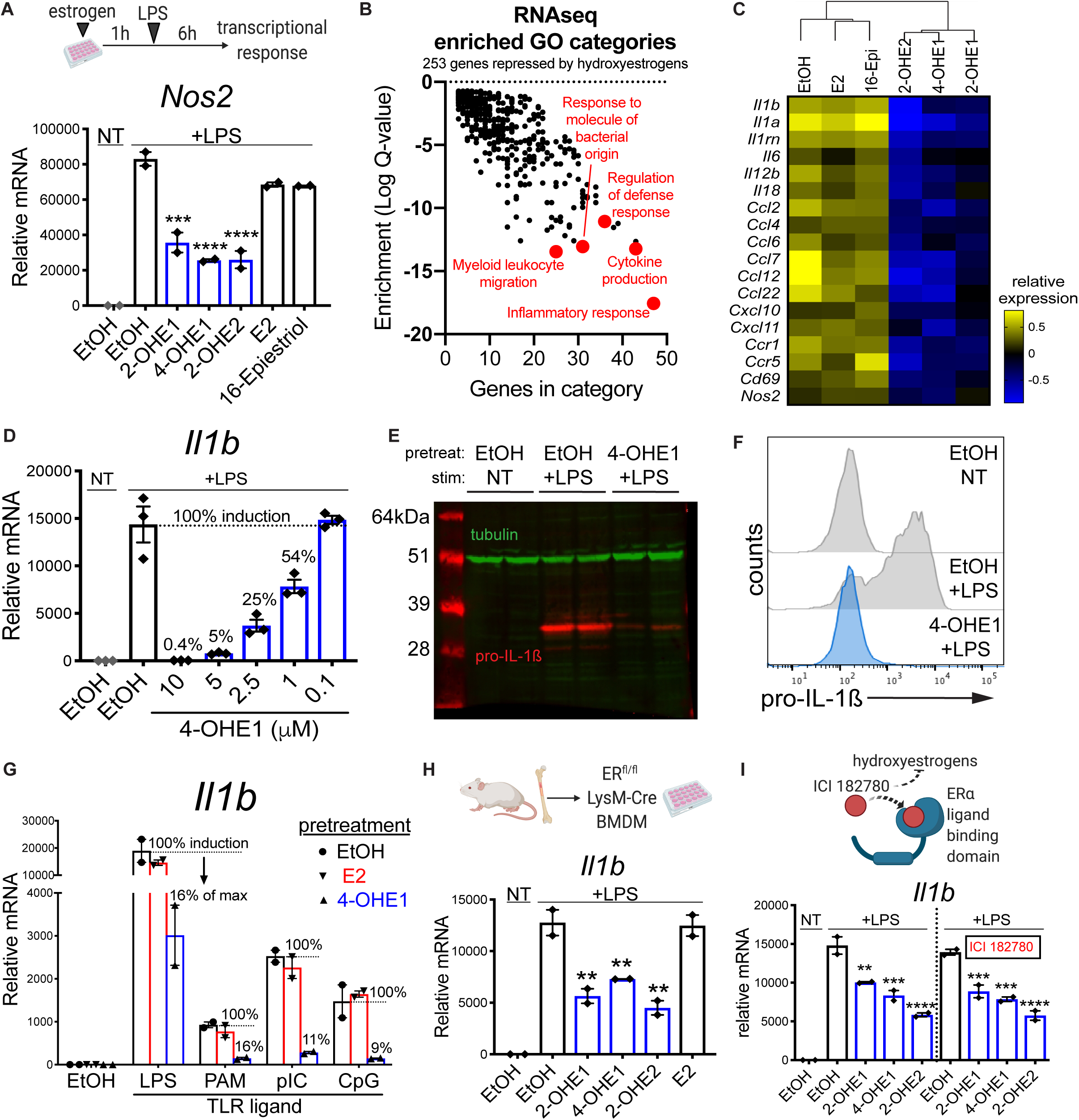
Hydroxyestrogens are anti-inflammatory in macrophages *in vitro*. **A**. BMDMs pretreated with EtOH vehicle control or 1μM estrogens for 1h, followed by 6h LPS stimulation (100ng/mL) and *Nos2* qPCR. **B**. Gene ontology (GO) analysis of 253 genes repressed by 1h hydroxyestrogen pretreatment (1μM) in 6h LPS-stimulated BMDMs identified by RNA-seq. **C**. RNA-seq hierarchical clustering dendrogram, with heatmap highlighting genes with reduced relative expression* in hydroxyestrogen-pretreated, LPS-stimulated BMDMs (*log2-transformed RPKM centered on the mean of each gene). **D**. BMDMs pretreated with EtOH or indicated concentrations of 4-OHE1 for 1h, followed by 6h LPS stimulation and *Il1b* qPCR. Percentages indicate induction relative to max (100%) in “EtOH +LPS” control BMDMs. **E**. RAW macrophages pretreated with EtOH or 5μM 4-OHE1 for 1h, followed by 6hLPS stimulation before pro-IL-1β measurement by western blot (n=2 independent biological replicates). **F**. RAW macrophages pretreated with EtOH or 5μM 4-OHE1 for 1h, followed by 4h LPS stimulation before pro-IL-1β measurement by intracellular staining and flow cytometry (representative data from 1 independent biological replicate for each condition shown). **G**. RAW macrophages pretreated with EtOH or 2.5μM estrogens for 1h, followed by 3h stimulation with LPS (TLR4), Pam3CSK4 (PAM, TLR2), polyinosinic-polycytidylic acid (pIC, TLR3), or CpG oligodeoxynucleotides (CpG, TLR9), and *Il1b* qPCR. Percentages indicate induction relative to max (100%) in the “EtOH +TLR ligand” control BMDMs for each ligand. **H**. BMDMs from ERα^fl/fl^ LysM-Cre mice pretreated with EtOH or 1μM estrogens for 1h, followed by 6h LPS stimulation (100ng/mL) and *Il1b* qPCR. **I**. BMDMs pretreated with EtOH or 1μM hydroxyestrogens in the absence (left) or presence (right) of 10μM ICI 182780 for 1h, followed by 6h LPS stimulation and *Il1b* qPCR. All qPCR data represented as mean ± SEM. Each data point is an independent biological replicate (n=2 or 3 for each condition). **P<0.01, ***P<0.001, ****P<0.0001 by one-way ANOVA with Dunnett’s test versus appropriate “EtOH +LPS” control sample. All qPCR data representative of at least 2 independent experiments.

**Supplementary Figure 1.**
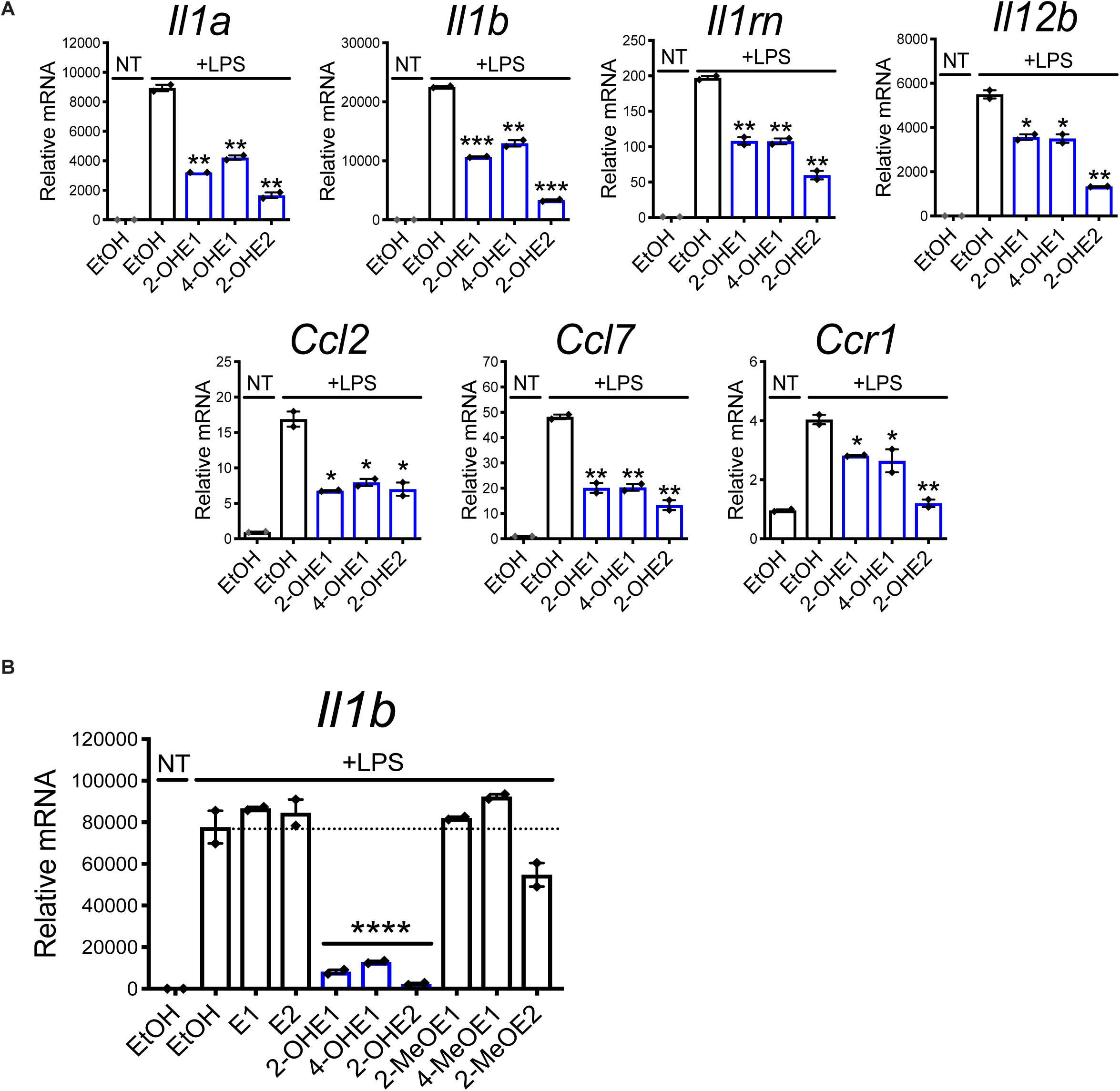
**A**. BMDMs pretreated for 1 hour with EtOH vehicle control or 1μM estrogens before 6 hour LPS stimulation (100ng/mL) and qPCR to validate cytokine and chemokine targets repressed by hydroxyestrogens from RNA-seq data.**B**. RAW macrophages pretreated for 1 hour with EtOH vehicle control or 5μM estrogens before 6 hour LPS stimulation (100ng/mL) and *Il1b* qPCR. Data represented as mean ± SEM. Each data point is an independent biological replicate (n=2 for each condition). *P<0.05, **P<0.01, ***P<0.001 by unpaired, two-sided Student’s T Test versus “EtOH +LPS” control sample (planned comparison) in **A**, or by one-way ANOVA with Dunnett’s test versus “EtOH +LPS” control sample in **B**.

**(Supplementary Fig.1A**). We chose *Il1b* as our model hydroxyestrogen-repressed gene as its transcriptional induction in BMDMs was potently repressed in a dose-dependent manner by 4-OHE1 pretreatment (**Fig.1D**). Strong *Il1b* repression by hydroxyestrogens also occurred in RAW macrophages (**Supplementary Fig.1B**), making them suitable for mechanistic studies. In agreement with the transcriptional effects, 4-OHE1 pretreatment strongly repressed LPS-induced pro-IL-1β protein levels (**Fig.1E,F**). 4-OHE1 pretreatment also repressed *Il1b* induction by multiple TLR agonists (**Fig.1G**), suggesting that a common pathway downstream of all TLRs is targeted.

To test if these anti-inflammatory effects were dependent on ERα, the primary ER expressed in macrophages^25, 26^, we repeated these experiments using BMDMs from ERα^fl/fl^ LysM-Cre mice. Surprisingly, the anti-inflammatory activity of the hydroxyestrogens was still intact (**Fig.1H**). Moreover, cotreatment with the high-affinity ERα antagonist ICI 182780 had no effect on the ability of the hydroxyestrogens to repress *Il1b* (**Fig.1I**). Together, these results demonstrate hydroxyestrogens are strong repressors of macrophage proinflammatory gene transcription *in vitro* that act in an ERα-independent manner.

### Hydroxyestrogens are anti-inflammatory *in vivo*

To test whether hydroxyestrogens have anti-inflammatory activity *in vivo*, we first examined 4-OHE1’s effects on acute LPS-induced inflammation. Mice were intraperitoneally (IP) injected with EtOH vehicle control, E2, or 4-OHE1, followed by IP injection with LPS (**Fig.2A**, top). 4-OHE1, but not E2, significantly repressed both the LPS-induced increase in serum IL-1β levels (**Fig.2A**, bottom), and LPS-induced proinflammatory gene expression in splenocytes (**Fig.2B**). Thus, 4-OHE1, but not E2, repressed acute LPS-induced inflammation *in vivo*.

**Figure 2.**
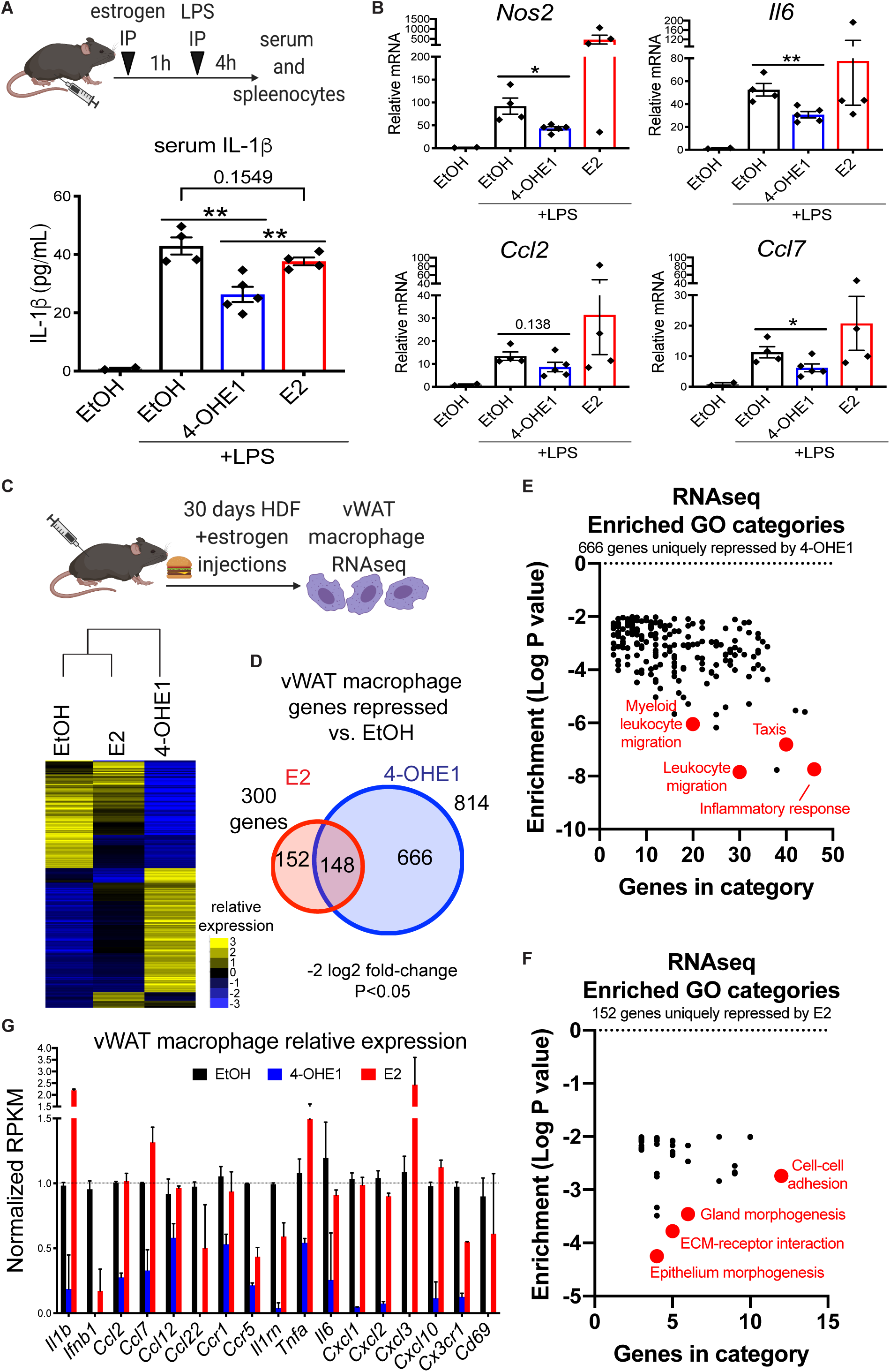
Hydroxyestrogens are anti-inflammatory *in vivo*. **A**. Experimental setup (top), and serum IL-1β levels (bottom) in 8-week-old C57BL/6 male mice injected IP with EtOH vehicle control or estrogens (10mg/kg) prior to IP LPS injection (2mg/kg). n=2,4,5, and 4 mice for the EtOH, EtOH +LPS, 4-OHE1 +LPS, and E2 +LPS groups, respectively. .**P<0.01 by unpaired, two-sided Student’s T Test (planned comparisons). **B**. qPCR for proinflammatory gene expression in splenocytes isolated from mice in **A**. *P<0.05, **P<0.01 by unpaired, two-sided Student’s T Test (planned comparisons). **C**. Experimental setup (top), and hierarchical clustering of vWAT macrophage RNA-seq data (bottom) from 8-week-old C57BL/6 male mice fed a high-fat diet (HFD) and injected subcutaneously every 6 days with EtOH, E2, or 4-OHE1 (10mg/kg). Relative expression heatmap displays the log2-transformed RPKM centered on the mean of each gene. n=5 mice for all groups. **D**. Venn Diagram displaying overlap between genes significantly repressed by E2 or 4-OHE1 versus EtOH control in vWAT macrophages from HFD-fed mice. **E. F**. GO analysis of 666 and 152 genes uniquely repressed by 4-OHE1 (top) or E2 (bottom), respectively, versus EtOH control vWAT macrophages. **G**. Relative expression of select proinflammatory genes in vWAT macrophages from HFD-fed mice injected with EtOH, 4-OHE1, or E2. All bar graphs represented as mean ± SEM.

**Supplementary Figure 2.**
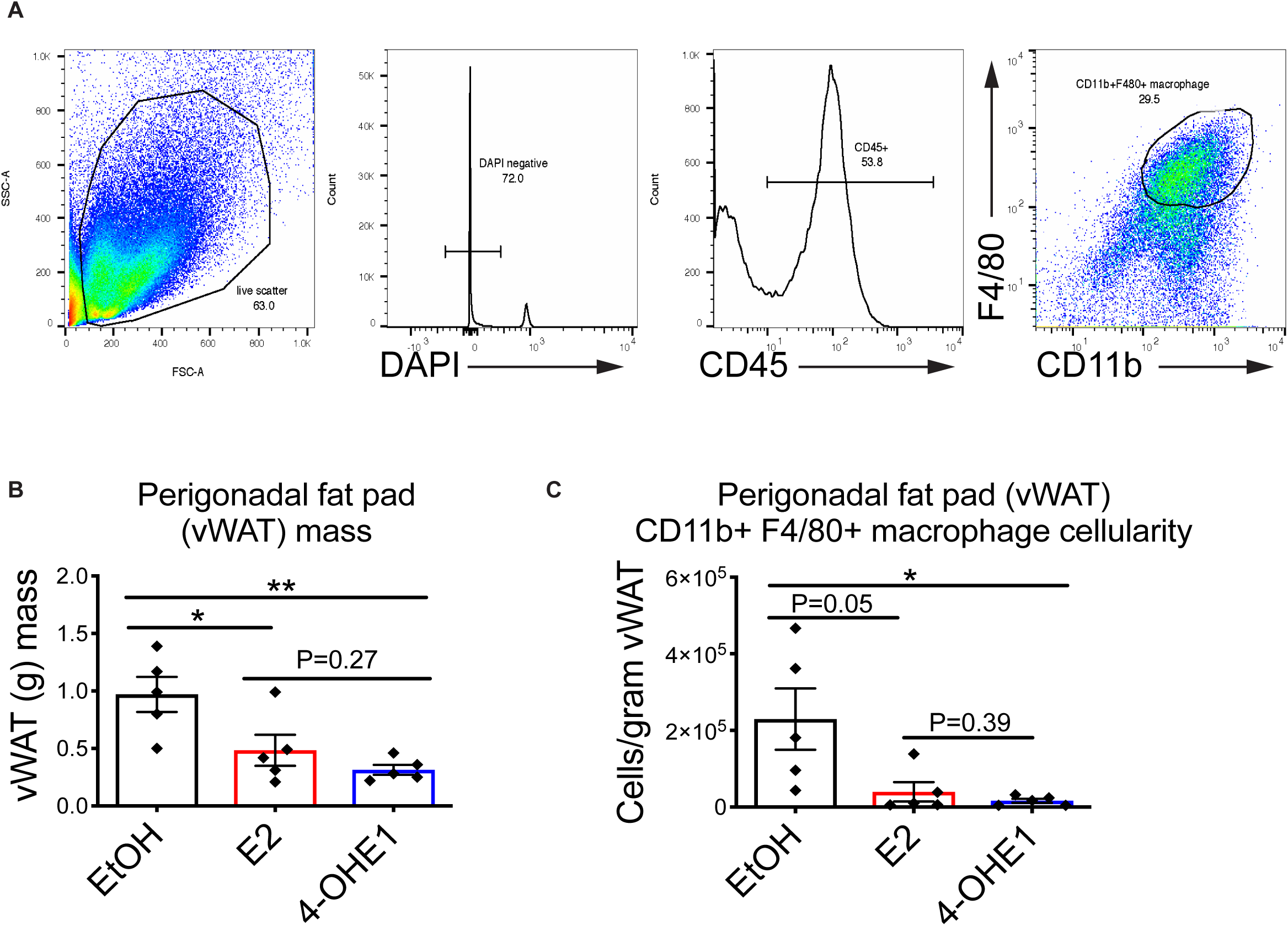
**A**. Representative gating strategy for identifying forward scatter/side scatter (FS/SS) live gate+, DAPI-, CD45+, F4/80+CD11b+ visceral white adipose tissue (vWAT) macrophages for sorting and flow cytometry analysis.**B**. vWAT mass in mice after 30 days HFD feeding and EtOH control or estrogen injection. *P<0.05, **P<0.01 by unpaired, two-sided Student’s T Test (planned comparisons).**C**. vWAT macrophage cellularity in mice after 30 days HFD feeding and EtOH control or estrogen injection. *P<0.05 by unpaired, two-sided Student’s T Test (planned comparisons). Bar graph data represented as mean ± SEM. n=5 mice per group.

To test if the hydroxyestrogens have anti-inflammatory effects *in vivo* in a chronic inflammatory setting, we examined effects on gene expression in visceral white adipose tissue (vWAT) macrophages during the early stages of high-fat diet (HFD)-induced metabolic inflammation. Mice were placed on HFD and given subcutaneous injections of EtOH, E2, or 4-OHE1 every 6 days. After 30 days, vWAT macrophages were profiled by RNA-seq (**Fig.2C**, top, **Supplementary Fig.2A**). While both E2 and 4-OHE1 reduced adiposity and macrophage cellularity in vWAT (**Supplementary Fig.2B,C**), vWAT macrophages from 4-OHE1-treated mice displayed a distinct gene expression profile compared to macrophages from EtOH and E2-treated mice (**Fig.2C**, bottom). 4-OHE1 repressed expression of a distinct set of genes compared to E2 (**Fig.2D, Supplementary Table 3**). GO analysis of genes uniquely repressed by 4-OHE1 revealed enrichment for categories including “Inflammatory response” and “Leukocyte migration”, (**Fig.2E**), while genes uniquely repressed by E2 showed no enrichment for inflammatory processes (**Fig.2F**). Many hydroxyestrogen targets repressed in macrophages *in vitro* were also repressed by 4-OHE1, but not E2, in vWAT macrophages (**Fig.2G**). Thus, 4-OHE1, but not E2, displays anti-inflammatory effects in vWAT macrophages during HFD-induced metabolic inflammation *in vivo*.

### Hydroxyestrogens activate NRF2, but NRF2 is dispensable for their anti-inflammatory activity

Given their ERα-independent anti-inflammatory effects, we considered other mechanisms by which hydroxyestrogens might act given what is known about their natural production and metabolism (**Fig.3A**). Hydroxylation of estrone (E1) or E2 by CYP1 family cytochrome P450 monooxygenases creates a catechol moiety, giving hydroxyestrogens their “catechol estrogens” nickname^27-29^. Present in a variety of drugs^30^ and natural compounds^31^, this catechol moiety can cause cellular stress in two ways. First, it can be oxidized to its quinone form, and redox cycling between these forms (which involves a semiquinone free radical intermediate) can produce reactive oxygen species (ROS)^32^. Second, because the quinone form possesses α,β-unsaturated carbonyls and is highly electrophilic, it can be attacked by nucleophiles, such as reactive cysteines on proteins, forming covalent adducts^33^. Accordingly, cells have evolved two ways to detoxify hydroxyestrogens: catechol methylation by COMT (which reduces redox cycling), and glutathione (GST) conjugation of the quinone. Given that hydroxyestrogens, but not their precursors or methylated metabolites, repressed LPS-induced proinflammatory gene transcription (**Fig.3B, Supplementary Fig.1B**), we hypothesized their anti-inflammatory activity must be dependent on their ability to cause oxidative and electrophilic stress.

**Figure 3.**
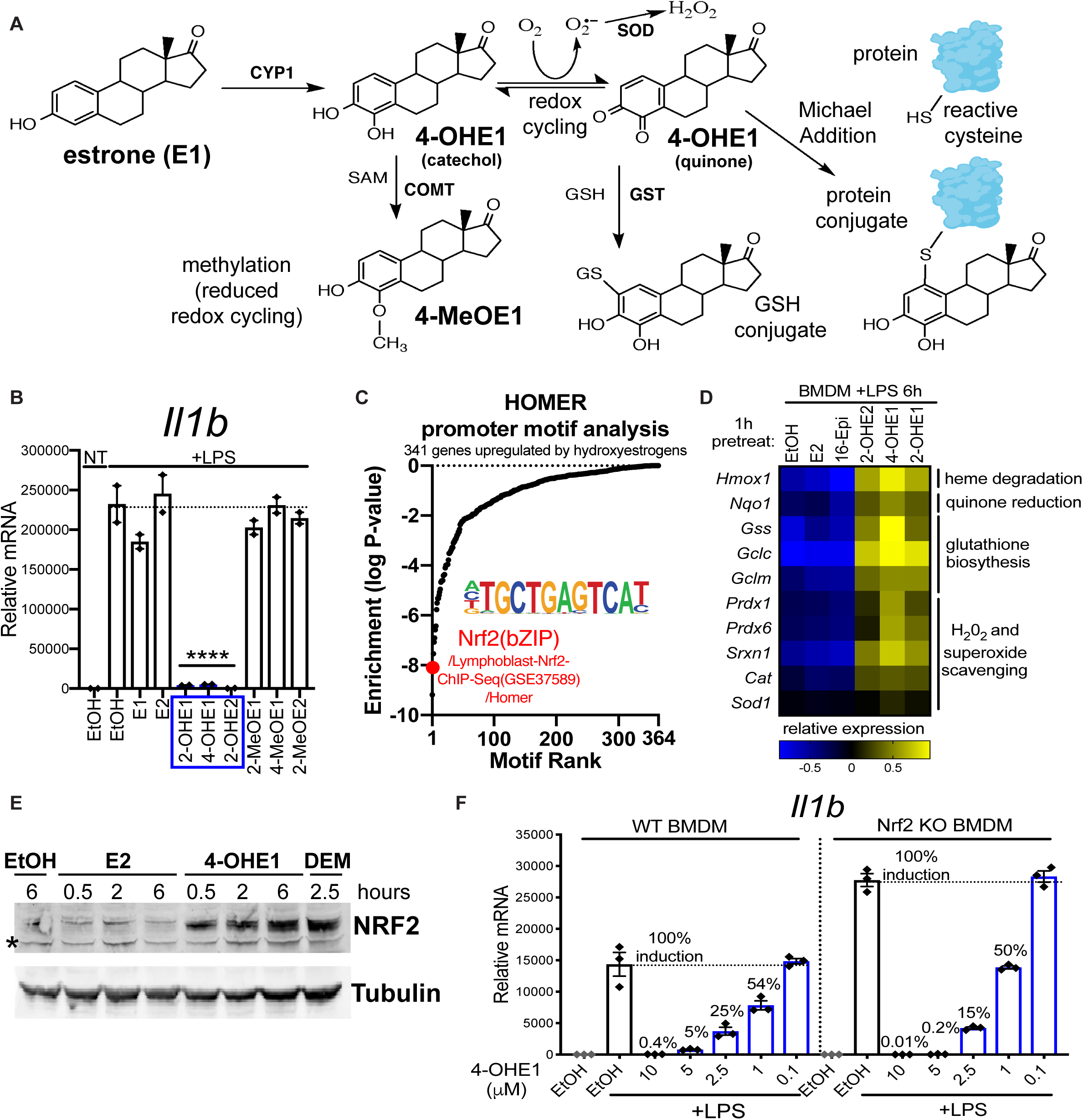
Hydroxyestrogens activate NRF2, but NRF2 is dispensable for their anti-inflammatory effects. **A**. Natural production and metabolism of 4-OHE1, highlighting detoxification via methylation and glutathione (GSH) conjugation, and mechanisms by which 4-OHE1 can cause oxidative and electrophilic stress. **B**.BMDMs pretreated with EtOH or 5μM estrogens for 1h, followed by 4h LPS stimulation and *Il1b* qPCR. ****P<0.0001 by one-way ANOVA with Dunnett’s test versus appropriate “EtOH +LPS” control sample. **C**. HOMER promoter motif analysis showing NRF2 as a top transcription factor binding motif enriched in promoters of the 341 genes upregulated by 1h hydroxyestrogen pretreatment (1μM) in 6h LPS-stimulated BMDMs. **D**.Relative expression* of putative NRF2 target genes in estrogen pretreated, LPS-stimulated BMDM RNA-seq dataset (*log2-transformed RPKM centered on the mean of each gene). **E**. RAW macrophages treated with 5μM E2, 5μM 4-OHE1, or 100μM DEM, for indicated times before NRF2 stabilization was assessed by western blot (*non-specific band). **F**.WT and Nrf2 KO BMDMs were pretreated 1h with EtOH or indicated concentrations of 4-OHE1 before 6h LPS stimulation and *Il1b* qPCR. Percentages indicate induction relative to max (100%) in “EtOH +LPS” control BMDMs for each genotype. WT BMDM data is from **Fig.1D**. **G**.qPCR data represented as mean ± SEM. Each data point is an independent biological replicate (n=2 or 3 for each condition). qPCR and western blot data representative of 2 independent experiments.

**Supplementary Figure 3.**
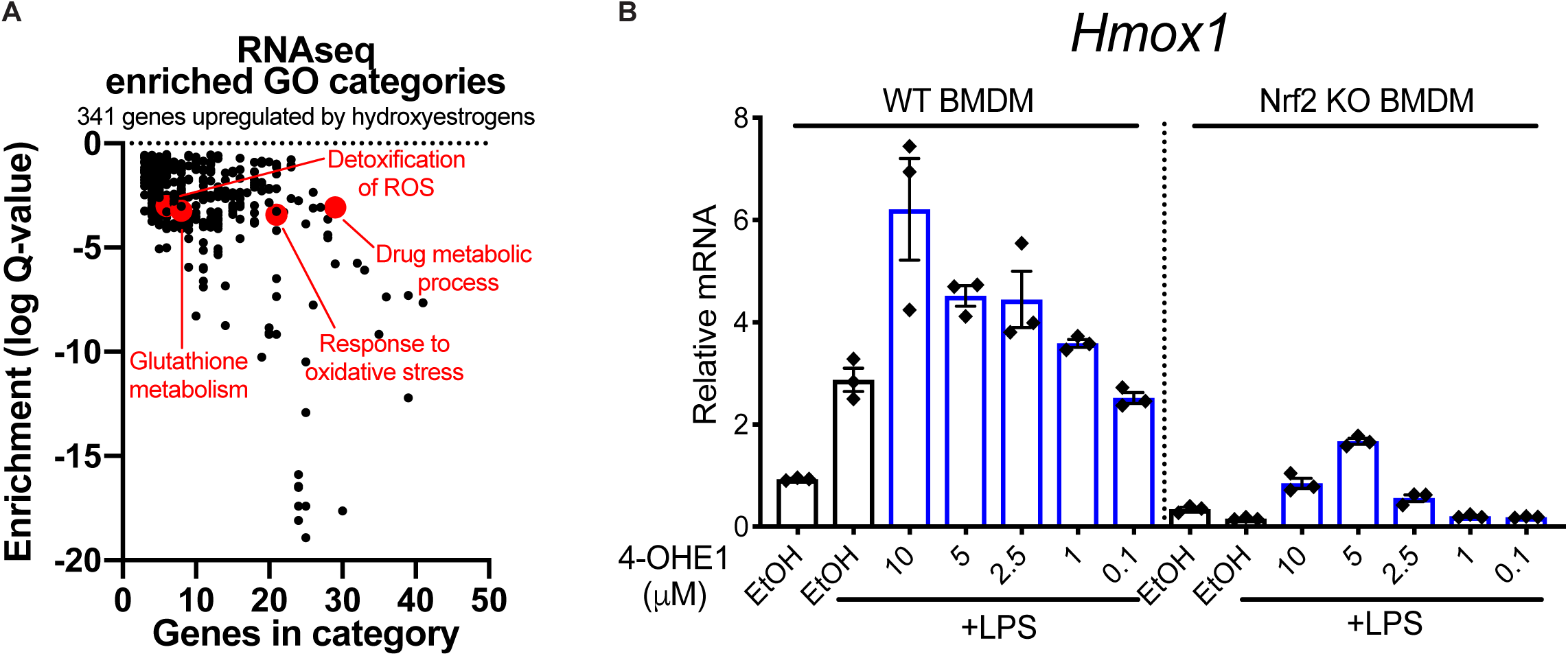
**A**.GO analysis of 341 genes significantly upregulated in hydroxyestrogen-pretreated, LPS-stimulated BMDMs relative to control pretreatments. Red highlights GO categories involved in oxidative stress resistance (OSR) and detoxification of reactive oxygen species (ROS). **B**. WT and Nrf2 KO BMDMs were pretreated 1h with EtOH or indicated concentrations of 4-OHE1 before 6h LPS stimulation (100ng/mL) and qPCR for the NRF2 target gene *Hmox1*. Data represented as mean ± SEM. n=3 independent biological replicates per condition.

The Keap1-Nrf2 system regulates cytoprotection in response to both oxidative and electrophilic stress. Our LPS-treated BMDM RNA-seq dataset identified 341 genes significantly upregulated in hydroxyestrogen-pretreated cells versus control pretreatments (**Supplementary Table 2**). GO analysis revealed enrichment of categories including “Response to oxidative stress”, “Detoxification of ROS”, and “Glutathione metabolism”, (**Supplementary Fig.3A**), and NRF2 was a top transcription factor binding motif enriched in promoters of these genes (**Fig.3C**). Expression of NRF2 targets *Hmox1, Nqo1*, and GST biosynthesis and ROS detoxification genes, were significantly upregulated by hydroxyestrogen pretreatment (**Fig.3D**). 4-OHE1 rapidly stabilized NRF2 in macrophages, similar to diethyl maleate (DEM), a known electrophilic NRF2 activator^34^ (**Fig.3E**). Given NRF2 has been described as a negative regulator of LPS-induced inflammation via its antioxidant activity^35^, and through direct repression of proinflammatory gene transcription^34^, we tested if NRF2 was required for 4-OHE1’s anti-inflammatory activity. BMDMs were prepared from wild-type (WT) and *Nfe2l2*^-/-^ (referred to as Nrf2 KO) mice. Impaired *Hmox1* induction confirmed lack of NRF2 function in Nrf2 KO BMDMs (**Supplementary Fig.3B**). However, 4-OHE1 repressed LPS-induced *Il1b* in both WT and Nrf2 KO BMDMs (**Fig.3F)**, demonstrating that while hydroxyestrogens are NRF2 activators, NRF2 is dispensable for their anti-inflammatory activity.

### Hydroxyestrogens cause mitochondrial stress

A recent report demonstrated E2 is highly abundant in mitochondrial membranes, and that E2 from distal and exogenous sources localizes to mitochondria in target cells and tissues^23^. Given NRF2 monitors mitochondrial integrity^36, 37^, we hypothesized NRF2 activation by hydroxyestrogens might be indicative of mitochondrial stress caused by these lipophilic compounds as they localize to mitochondria, producing ROS and covalently modifying mitochondrial proteins. GO analysis of the 341 genes upregulated in hydroxyestrogen-pretreated LPS-stimulated BMDMs revealed three additional transcriptional signatures indicative of mitochondrial stress (**Fig.4A-C, Supplementary Fig.4A-C**.). The Heat Shock Factor 1 (HSF1) signature includes putative HSF1 target genes^38-40^, suggesting HSF1 activation in response to mitochondrial dysfunction^41-46^. The ATF4/mitochondrial damage signature includes *Atf4*, which coordinates cytoprotection in response to mitochondrial stress in mammalian cells^47^, and the ATF4 target *Gdf15*^48^, encoding a mitokine indicative of mitochondrial dysfunction^49^. Finally, the Glycolysis/Pentose Phosphate Pathway (PPP) signature includes enzymes in these pathways known to be upregulated in response to mitochondrial stress in other systems^16, 50^.

**Figure 4.**
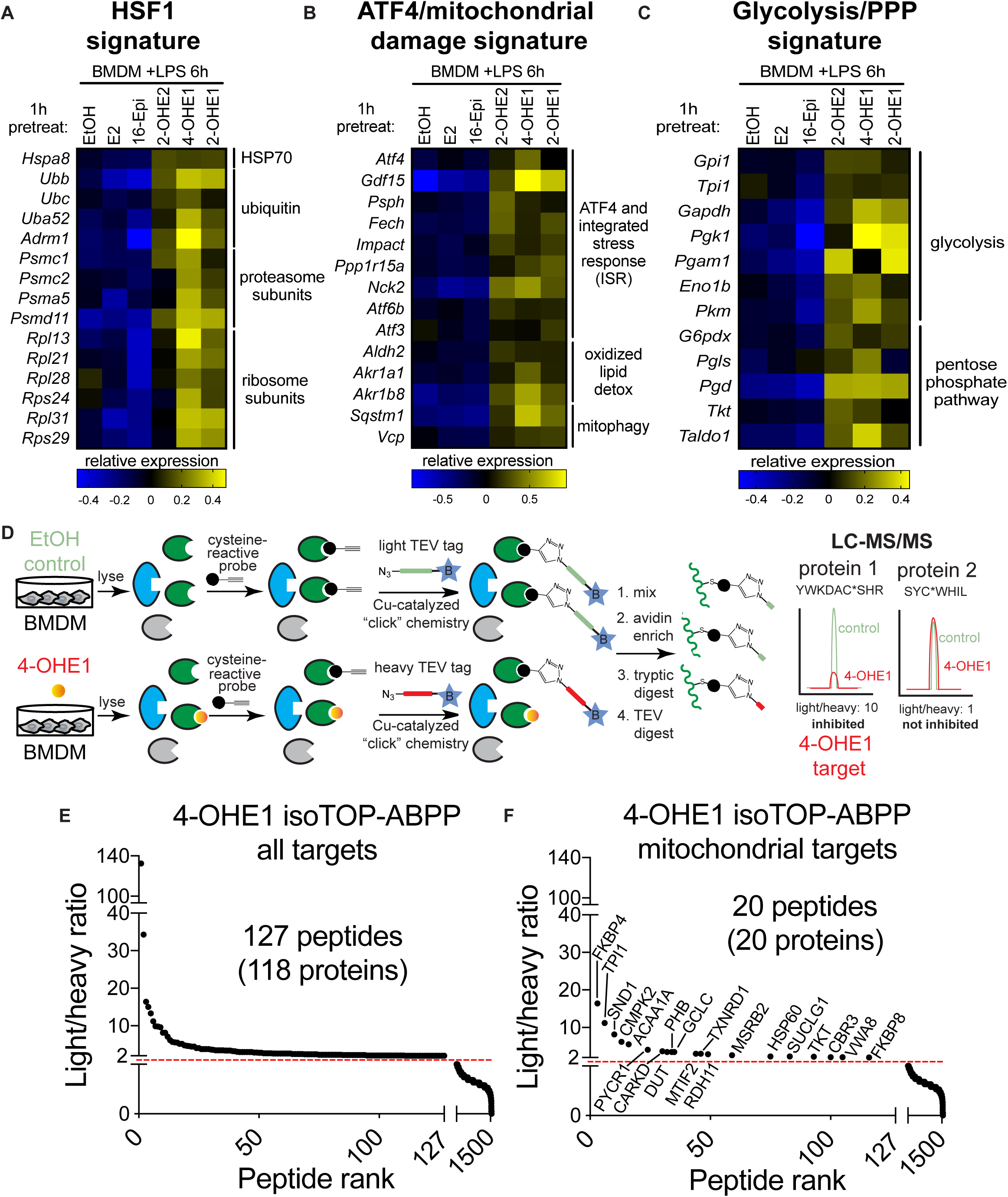
Hydroxyestrogens cause mitochondrial stress. **A.B.C**. Relative expression* of genes indicative of mitochondrial stress in 1h estrogen pretreated, 6h LPS-stimulated BMDM RNA-seq dataset (*log2-transformed RPKM centered on the mean of each gene). **D**. IsoTOP-ABBP strategy to identify covalent targets of 4-OHE1 acting through reactive cysteines. BMDMs were treated with EtOH or 1μM 4-OHE1 for 1h (n=3 independent biological replicates per condition). **E. F**. All targets (left) and mitochondrial targets (right) of 4-OHE1 identified by isoTOP-ABPP. Cysteine-containing peptides above dashed line have light/heavy ratio>2.0, indicating at least a 50% reduction in cysteine-reactive probe targeting of these cysteines in 4-OHE1-treated BMDMs relative to control BMDMs. 18 of 20 mitochondrial targets are in MitoCarta 2.0. FKBP4 and GCLC mitochondrial localization prediction from UniProt.

**Supplementary Figure 4.**
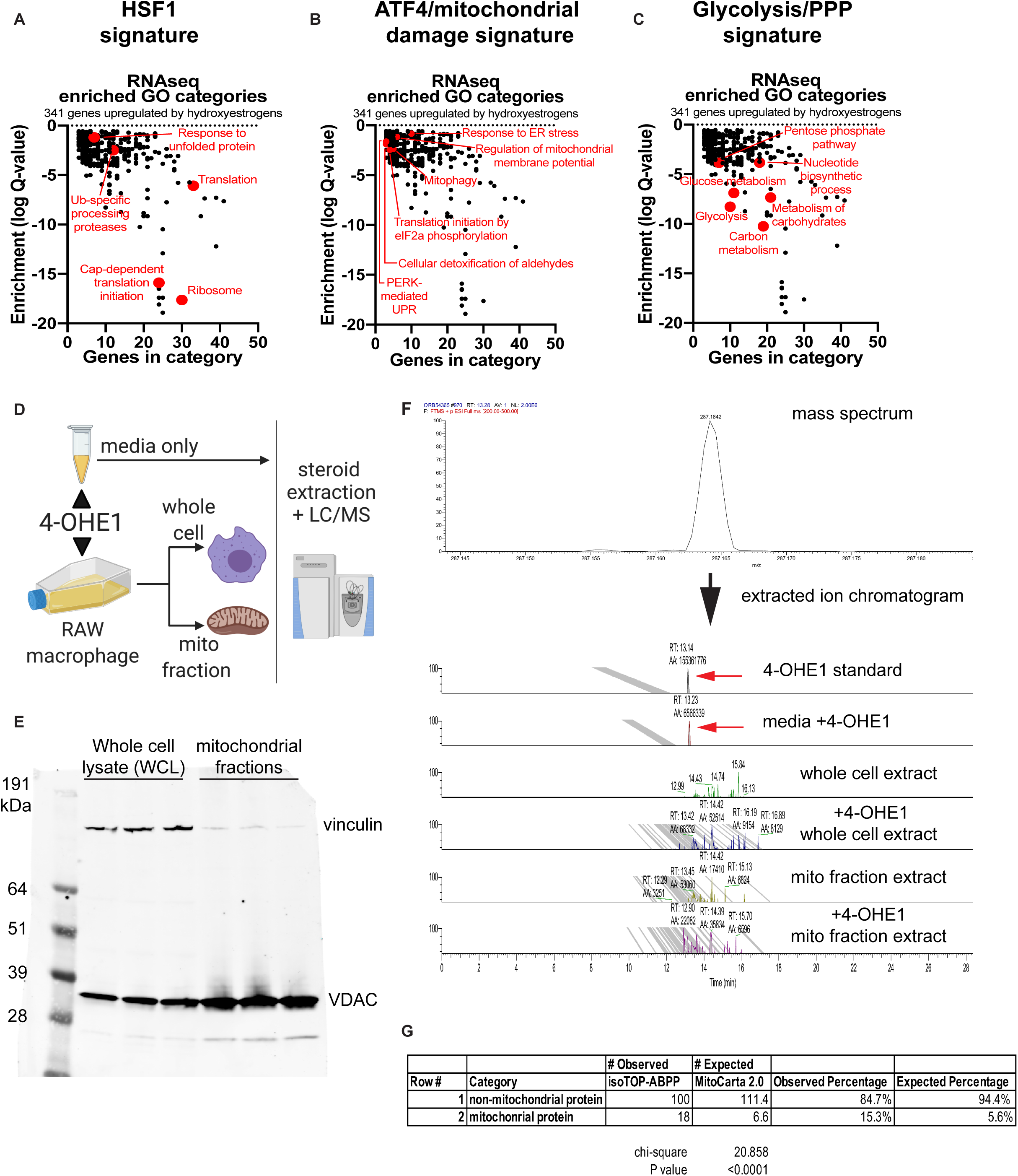
**A-C**. GO analysis of 341 genes significantly upregulated in hydroxyestrogen-pretreated, LPS-stimulated BMDMs relative to control pretreatments. Red highlights GO categories indicative of HSF1 and ATF4 activity, and upregulation of glycolysis/pentose phosphate pathway (PPP) genes.**D**. Experimental setup for steroid extraction and liquid chromatography/mass spectrometry (LC-MS) to measure 4-OHE1 extracted from cell culture media (top, control), or from whole cell and mitochondrial fractions prepared from RAW macrophages treated with 5μM 4-OHE1 for 1 hour (n=2 independent biological replicates per condition). **E**.Representative western blot confirming enrichment of mitochondrial marker VDAC, and depletion of cytoplasmic marker vinculin, in mitochondrial fractions versus whole cell lysates prepared from RAW macrophages. **F. top** – Mass spectrum (positive ion mode) measured for 1mg/mL 4-OHE1 standard (injection volume = 2μL) showing detail for the [M+H]^+^ ion of 4-OHE1 at *m/z* = 287.1642. Mass range displayed: *m/z* = 287.1624-287.1660 **bottom** – Extracted ion chromatograms for 4-OHE1 (*m*/*z* = 287.1642). Peaks at retention time (RT) = 13.14 and RT=13.23 minutes for the 4-OHE1 standard and media +4-OHE1 sample (denoted by red arrows) confirm the ability of our LC-MS method to detect 4-OHE1, and validates our 4-OHE1 steroid extraction method. Lack of a defined chromatographic peak at RT = 13.1-13.2 minutes for whole cell or mitochondrial extracts prepared from RAW macrophages treated with 4-OHE1 indicates lack of a detectable quantity of free 4-OHE1 in these samples. Data is representative of that obtained with n=2 independent biological replicates per condition. **G**.Chi-square test comparing the observed frequency of mitochondrial targets (18) in our isoTOP-ABPP target list (118 total targets) versus the expected frequency of mitochondrial targets from MitoCarta 2.0.

We performed steroid extraction^51^ and liquid chromatography/mass spectrometry (LC-MS) to determine if 4-OHE1 was enriched in mitochondrial fractions relative to whole cells (**Supplementary Fig.4D,E**). However, we were unable to detect free 4-OHE1 in macrophages treated with 4-OHE1 for 1 hour (**Supplementary Fig.4F**), suggesting 4-OHE1 is rapidly covalently conjugated by GST and/or proteins (**Fig.3A**). To identify covalent 4-OHE1 protein targets acting through reactive cysteines, we performed competitive isotopic tandem orthogonal proteolysis-enabled activity-based protein profiling (isoTOP-ABPP)^52^ in EtOH control and 4-OHE1-treated BMDMs (**Fig.4D**). This revealed 127 cysteines on 118 proteins targeted by 4-OHE1 (**Fig.4E**). Cross-referencing this target list with MitoCarta 2.0^53^ revealed 18 of the 118 targets are mitochondrial proteins (**Fig.4F**). A chi-square test comparing the observed frequency of mitochondrial targets with the expected frequency of mitochondrial proteins from MitoCarta 2.0 confirmed significant enrichment for mitochondrial proteins in the 4-OHE1 target list (**Supplementary Fig.4G**). Together, these transcriptional signatures and proteomic data suggest hydroxyestrogens cause mitochondrial stress via local ROS production and covalently targeting mitochondrial proteins.

### Hydroxyestrogens impair mitochondrial acetyl-CoA production and histone acetylation required for LPS-induced proinflammatory gene transcription

We next considered how oxidative and electrophilic mitochondrial stress caused by hydroxyestrogens might exert anti-inflammatory effects. Mitochondria utilize oxidized glucose for acetyl-CoA production, which in turn is used to acetylate histones^54, 55^, a process crucial for *Il1b* upregulation in LPS-stimulated macrophages^56, 57^.

Pharmacological inhibition of glucose utilization in LPS-stimulated macrophages has selective repressive effects on *Il1b*, but not cytokine transcripts such as *Il6* and *Tnf*, suggesting the glucose-to-acetyl-CoA axis is more important for transcription of the former gene^58, 59^. Similarly, in both RAW macrophages (**Fig.5A**) and BMDMs (**Supplementary Fig.5A**), 4-OHE1 pretreatment had much stronger repressive effects on LPS-induced *Il1b* than *Il6* and *Tnf*, suggesting 4-OHE1 interferes with mitochondrial glucose utilization for acetyl-CoA production.

**Figure 5.**
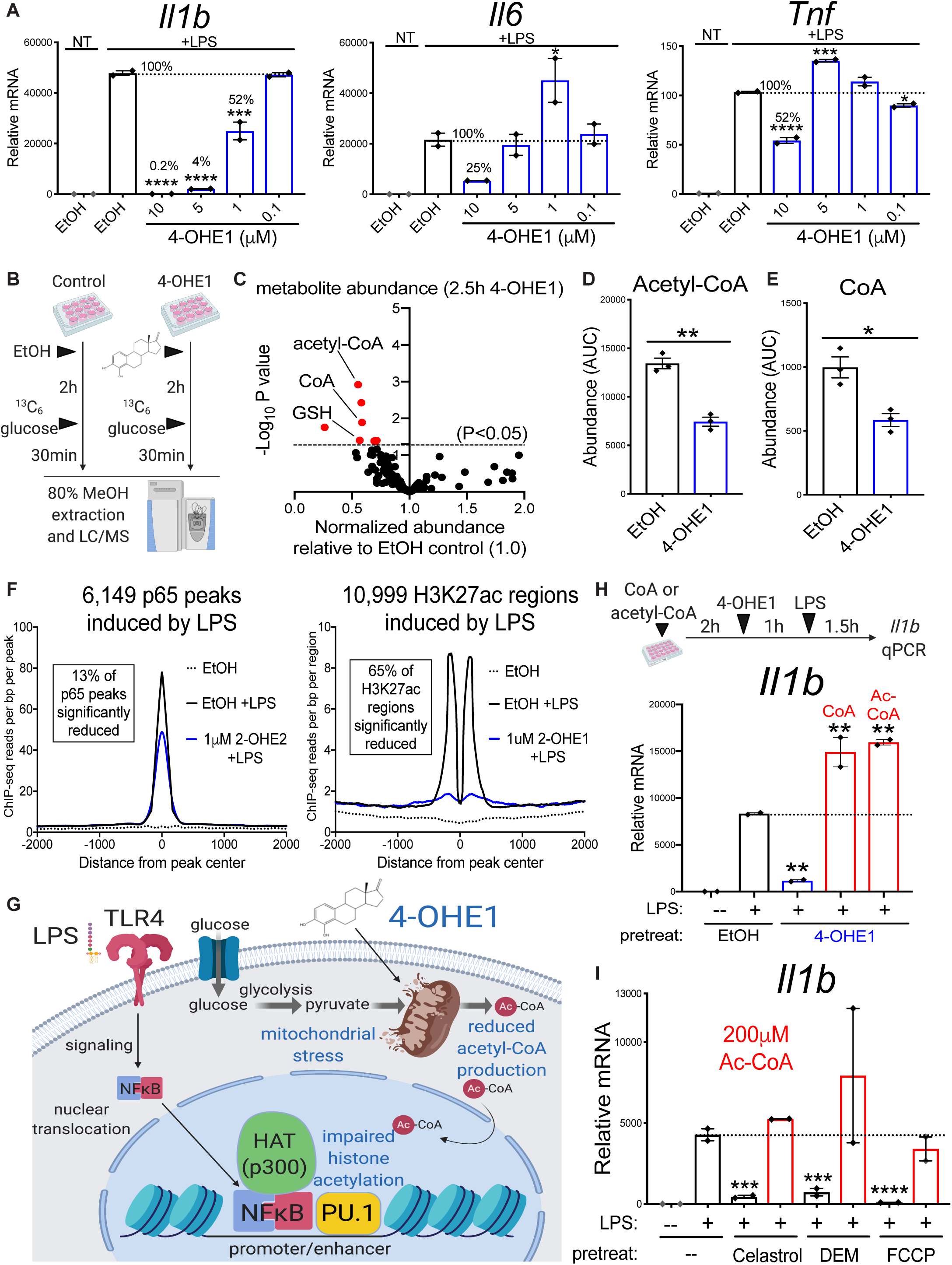
Hydroxyestrogens impair mitochondrial acetyl-CoA production and histone acetylation required for LPS-induced proinflammatory gene transcription. **A**. RAW macrophages pretreated for 1h with indicated concentrations 4-OHE1 before LPS stimulation for 6h and *Il1b, Il6*, and *Tnf* qPCR. **B**. BMDM metabolomics and ^13^C6-glucose labeling strategy (n=3 independent biological replicates per condition) **C**.Normalized metabolite abundance in 4-OHE1-treated BMDMs (5μM) relative to EtOH control BMDMs (where metabolite level was set to 1.0). Red highlighted metabolites above dashed line showed significant change (one-way ANOVA, P<0.05). **D. E**. Absolute abundance of acetyl-CoA (left) and CoA (right) in EtOH and 4-OHE1-treated BMDMs. *P<0.05, **P<0.01 by unpaired, two-sided Student’s T test. **F.** ChIP-seq for p65 (left) and H3K27ac (right) in BMDMs pretreated for 1h with 1μM of the indicated hydroxyestrogen, followed by 30 minute LPS stimulation. Histograms represent read density at LPS-inducible p65 peaks/H3K27ac regions in each of the 3 conditions. H3K27ac regions were centered on the nucleosome-free region (NRF) of associated promoters/enhancers, yielding a classic “2 peak” histogram. **G.** Model for how mitochondrial stress caused by 4-OHE1 impairs metabolic/epigenetic control of proinflammatory gene transcription. **H.** BMDMs cultured in absence or presence of CoA (250μM) or acetyl-CoA (Ac-CoA, 200μM) (red bars) for 3h prior to 1h 5μM 4-OHE1 pretreatment, 1.5h LPS stimulation, and *Il1b* qPCR. **I.** RAW macrophages cultured in absence or presence of acetyl-CoA (Ac-CoA, 200μM, red bars) for 2h prior to 1h electrophile pretreatment, 1.5h LPS stimulation, and *Il1b* qPCR. Concentrations: 250nM Celastrol, 100μM DEM, 10μM FCCP. All bar graphs represented as mean ± SEM, and each data point is an independent biological replicate (n=2 or 3 for each condition). For qPCR, *P<0.05, **P<0.01,***P<0.001, ****P<0.001 by one-way ANOVA with Dunnett’s test versus appropriate “EtOH +LPS” control sample. All qPCR data representative of at least 2 independent experiments.

**Supplementary Figure 5.**
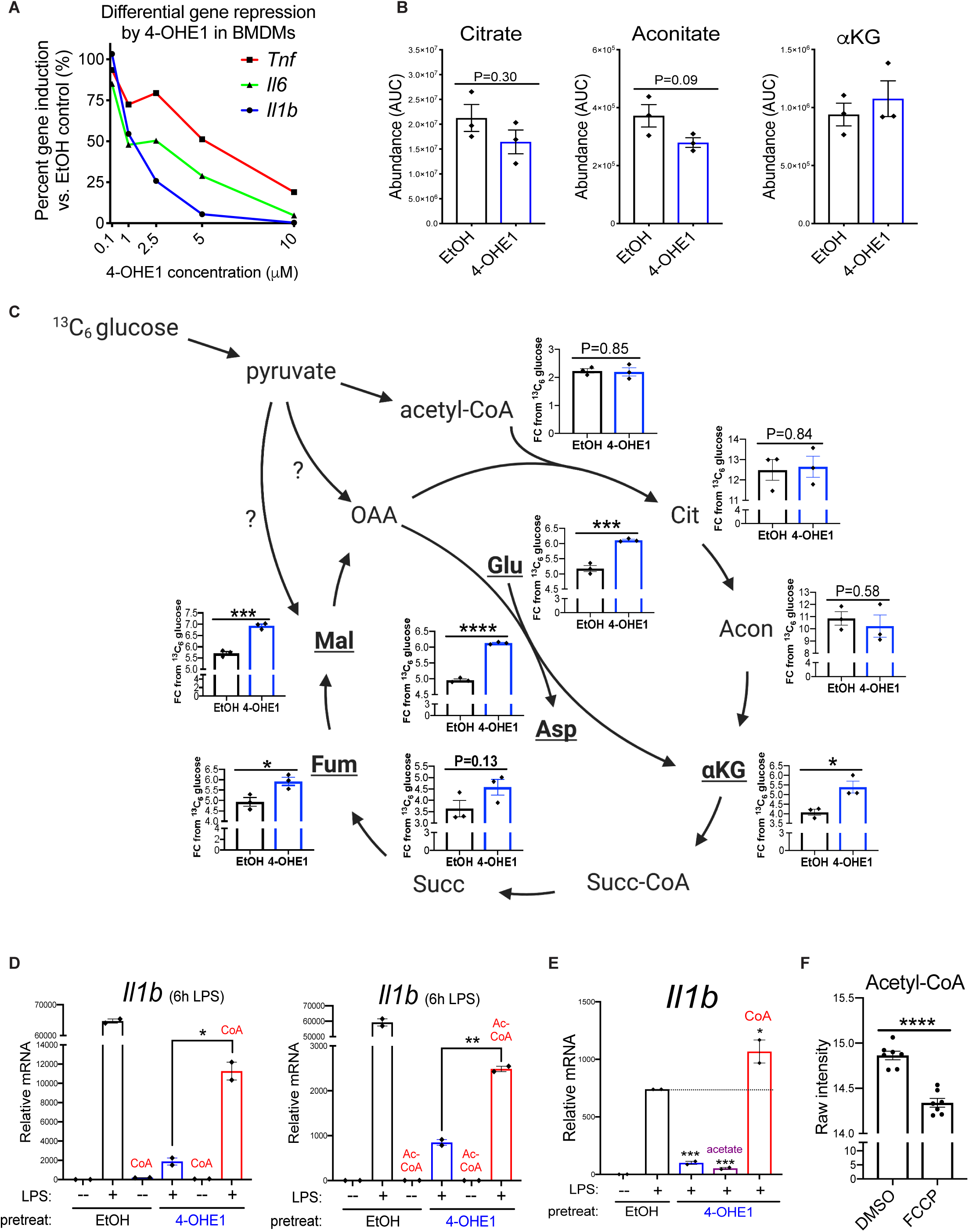
**A.** BMDMs were pretreated 1h with EtOH or indicated concentrations of 4-OHE1 before 6h LPS stimulation (100ng/mL) and qPCR for *Il1b, Il6*, and *Tnf*. The average percent induction of at each 4-OHE1 concentration ((induction in presence of 4-OHE1/EtOH control induction)x100)) was calculated from n=3 independent biological replicates for each data point and plotted for each gene. **B.** Absolute abundance (area under the curve, AUC) of citrate, aconitate, and alpha-ketoglutarate in EtOH versus 4-OHE1-treated BMDMs. P values from unpaied, two-sided Student’s T Test. **C.** Fractional contribution (FC) of ^13^C6-glucose-derived carbons to total carbons for TCA cycle metabolites, and amino acids derived from these metabolites, in EtOH versus 4-OHE1-treated BMDMs. Metabolites with significantly enhanced ^13^C labeling in bold. Question marks indicate possible alternative entry routes for ^13^C6-glucose-derived carbons into the TCA cycle other than pyruvate conversion to acetyl-CoA and citrate.*P<0.05, ***P<0.001, ****P<0.0001 by unpaired, two-sided Student’s T Test versus EtOH. **D**. RAW macrophages cultured in absence or presence of CoA (500μM, left) or acetyl-CoA (Ac-CoA, 500μM, right) (red bars) for 3 hours prior to 1 hour EtOH or 5μM 4-OHE1 pretreatment, 6 hour LPS stimulation (100ng/mL), and *Il1b* qPCR. *P<0.05, **P<0.01 by unpaired, two-sided Student’s T Test (planned comparison). **E**. RAW macrophages cultured in absence or presence of sodium acetate (5mM, purple bar) or CoA (500μM, red bar) for 15 minutes prior to 1 hour EtOH or 5μM 4-OHE1 pretreatment, 1 hour LPS stimulation (100ng/mL), and *Il1b* qPCR. *P<0.05, ***P<0.001 by one-way ANOVA with Dunnett’s test versus “EtOH +LPS” control sample. All bar graph data represented as mean ± SEM. n=2 or 3 independent biological replicates per condition. **F.** Data from reference 38. Acetyl-CoA levels as measured by mass spectrometry in HeLa cells treated for 24 hours with DMSO vehicle control or 10μM FCCP. Data represented as mean ± SEM, n=7 independent biological replicates for each groups.

To test this hypothesis, we performed a metabolomics screen to identify metabolites whose levels were acutely affected by 4-OHE1 treatment. BMDMs were treated with EtOH or 4-OHE1 for two hours, followed by 30 minute ^13^C6-glucose labeling before whole cell metabolite extraction (**Fig.5B**). Of the 136 metabolites measured, just eight were significantly changed by 4-OHE1 (**Fig.5C**). Free GSH levels were reduced (**Fig.5C**), supporting the notion that 4-OHE1 is rapidly glutathione-conjugated after treatment (**Fig.3A**). And in line with our hypothesis, the metabolite with the most statistically significant change was acetyl-CoA, as levels in 4-OHE1-treated BMDMs decreased by 45% (**Fig.5C,D**). Interestingly, Coenzyme A (CoA) levels were also significantly reduced (41%) by 4-OHE1 treatment (**Fig.5C,E**). As CoA biosynthesis occurs within mitochondria^60, 61^, and 80-90% of intracellular CoA is intra-mitochondrial^62, 63^, this suggests mitochondrial stress caused by 4-OHE1 disrupts CoA homeostasis, and in turn, acetyl-CoA production. For TCA cycle metabolites quantified, total levels of citrate and aconitate (immediately downstream of acetyl-CoA) showed the largest decreases in abundance (**Supplementary Fig.5B**). Moreover, our ^13^C6-glucose labeling revealed increased ^13^C fractional contribution to all TCA cycle metabolites (and amino acids derived from these metabolites) except citrate and aconitate, upon 4-OHE1 treatment (**Supplementary Fig.5C**), suggesting alternative entry routes of glucose-derived carbon into the TCA cycle other than via acetyl-CoA (also not increased) to replenish intermediates in response to 4-OHE1-induced mitochondrial stress^64^. Together, this data demonstrates mitochondrial stress caused by 4-OHE1 impairs mitochondrial acetyl-CoA production.

Since acetyl-CoA is required for histone acetylation and proinflammatory gene transcription, we performed ChIP-seq to take an unbiased, genome-wide look at how LPS-induced transcription factor binding and histone acetylation were affected by hydroxyestrogen pretreatment. For the NF*K*B subunit p65, we identified 6,149 peaks absent in unstimulated BDMDs but present 30 minutes post-LPS stimulation (**Fig.5F**, left). Pretreatment with the hydroxyestrogen 2-OHE2 had minimal effects on LPS-induced NF*ĸ*B binding, as only 13% of LPS-induced p65 binding peaks were significantly reduced in read density. For histone acetylation, we identified 10,999 regions of H3K27ac, a mark of active promoters and enhancers, absent in naïve BMDMs but present 30 minutes post-LPS stimulation (**Fig.5F**, right). In contrast with NF*ĸ*B binding, pretreatment with the hydroxyestrogen 2-OHE1 strongly impaired H2K27ac deposition, as nearly two-thirds (65%) of LPS-induced H3K27ac regions were significantly reduced in read density. Thus, while hydroxyestrogens largely leave TLR4 signaling and transcription factor nuclear translocation/DNA binding intact, they strongly impair histone acetylation required for proinflammatory gene transcription, further supporting the hypothesis that hydroxyestrogens impair mitochondrial acetyl-CoA production (**Fig.5G**).

Administration of exogenous CoA to cells with impaired CoA biosynthesis^65^, or treated with CoA-depleting compounds^66^, can rescue histone acetylation and gene expression defects caused by reduced acetyl-CoA levels. Indeed, supplying BMDMs with either exogenous CoA or acetyl-CoA fully rescued LPS-induced *Il1b* in 4-OHE1-pretreated macrophages at early timepoints (1-1.5h) post-LPS (**Fig.5H)**, and partially at 6 hours (**Supplementary Fig.5D**). Acetate supplementation was unable to rescue *Il1b* (**Supplementary Fig.5E**), suggesting nucleocytosolic acetyl-CoA generation by ASCC2 does not contribute to the acetyl-CoA pool in macrophages^67^. These rescue experiments further support a model where impairment of mitochondrial acetyl-CoA production by hydroxyestrogens underlies their anti-inflammatory activity.

Finally, we wondered if this anti-inflammatory mechanism might extend to other electrophilic small molecules with immunomodulatory properties^68, 69^. Celastrol and DEM are electrophiles that repress *Il1b* transcription in myeloid cells^34, 70^, and activate the same stress response pathways as 4-OHE1^71-74^. Mitochondrial uncouplers including carbonyl cyanide 4-(trifluoromethoxy)phenylhydrazone (FCCP) are known for their ability to dissipate mitochondrial membrane potential (mtMP) and reduce mtROS levels, an effect thought to be behind *Il1b* repression in macrophages by uncouplers as mtROS is considered an inflammatory signal^75^. However, uncouplers are also electrophilic^76-79^, and like 4-OHE1, FCCP treatment significantly reduces intracellular acetyl-CoA levels^47^ (**Supplementary Fig.5F**). Thus, we tested if acetyl-CoA supplementation could rescue LPS-induced *Il1b* expression in macrophages pretreated with these electrophiles. Indeed, supplying acetyl-CoA prior to electrophile treatment restored *Il1b* induction (**Fig.5I**), suggesting the anti-inflammatory effects of these electrophiles may also lie in their ability to cause mitochondrial stress and impair acetyl-CoA production.

### Hydroxyestrogen-driven mitochondrial stress triggers mitohormesis

We next wondered how the mitochondrial stress caused by hydroxyestrogens might influence macrophage function beyond these acute anti-inflammatory effects. High levels of mitochondrial stress trigger apoptosis; however, we observed no changes in mitochondrial oxygen consumption (**Supplementary Fig.6A**), mtMP (**Supplementary Fig.6B**), or cell viability upon acute hydroxyestrogen treatment. Alternatively, transient mild doses of mitochondrial stress can trigger persistent stress adaptations that provide cyto-/mito-protection during subsequent stress exposure in a process known as mitohormesis^17, 18^. Thus, we tested if hydroxyestrogen-driven mitochondrial stress triggered mitohormesis in macrophages.

**Figure 6.**
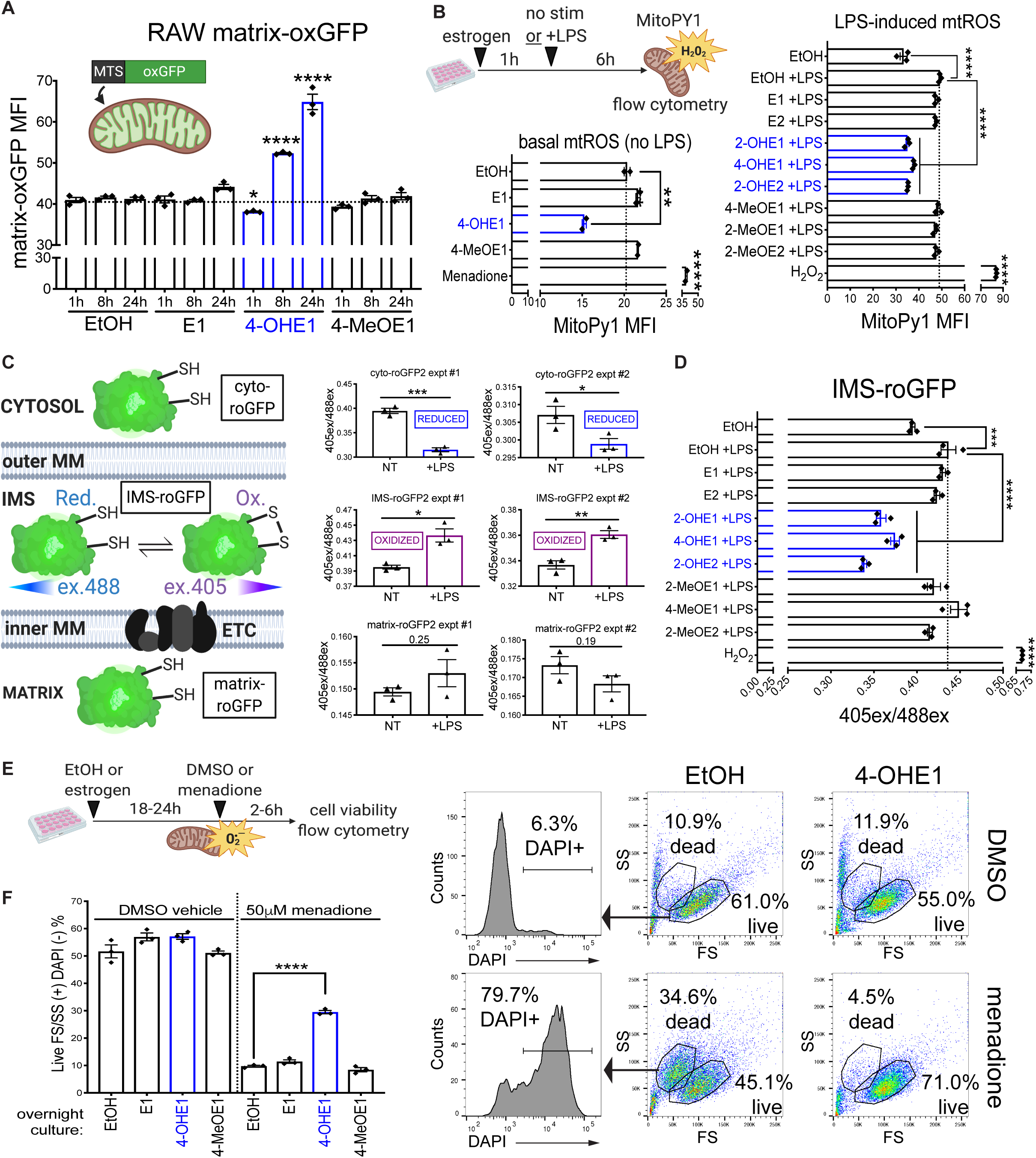
Hydroxyestrogen-driven mitochondrial stress triggers mitohormesis. **A**. RAW matrix-oxGFP macrophages were treated with EtOH or 5μM estrogens for indicated times and matrix-oxGFP florescence quantified by flow cytometry.**B**. Basal (left) and LPS-induced (right) mitochondrial H2O2 levels were measured in EtOH and estrogen (pre)treated (5μM) RAW macrophages using MitoPy1 staining and flow cytometry. Menadione (50μM) and H2O2 (500μM) serve as positive controls.**C. left** – Schematic describing RAW macrophages expressing roGFP proteins targeted to cytosol, mitochondrial inner membrane space (IMS), and mitochondrial matrix. roGFP oxidation favors excitation by 405nm violet laser relative to 488nm blue laser.**right** – 405nm/488nm excitation ratio measured by flow cytometry (510nM emission) in roGFP RAW macrophages untreated (NT) or treated with LPS (100ng/mL) for 6 hours.*P<0.05, **P<0.01, ***P<0.001, by unpaired, two-sided Student’s T Test.**D**. IMS-roGFP macrophages pretreated with EtOH or estrogens (5μM) for 1h and stimulated with LPS for 6h before 405/488nm excitation ratio was measured by flow cytometry (510nM emission).**E. left** – Schematic describing menadione mitochondrial superoxide resistance assay. RAW macrophages were treated overnight (18-24h) with EtOH or 4-OHE1 (5μM) before next day treatment with DMSO vehicle control or menadione (50μM, 4h) and viability assessment by flow cytometry. **right** – Flow cytometry forward scatter/side scatter plots (FS/SS) of treated RAW macrophages. DAPI staining demonstrates cells in “live” FS/SS gate are viable and exclude DAPI, whereas “dead” FS/SS gate cells have increased cell membrane permeability and take up DAPI. Percentages represent events in “live” and “dead” FS/SS gates relative to total events.**A**. RAW macrophages treated overnight with EtOH or 5μM estrogens were treated the following day with DMSO vehicle control (left) or 50μM menadione for 4h before viability was assessed the next day by flow cytometry. Viability is represented as percentage of “live” FS/SS gate positive, DAPI negative cells per total events collected for each sample. All flow cytometry data represented as mean ± SEM. Each data point is an independent biological replicate (n=2 or 3 for each condition). *P<0.05, ***P<0.001, and ****P<0.0001 by one-way ANOVA with Dunnett’s test versus appropriate “EtOH” control timepoint in **A**., or the appropriate control sample indicated by bars/arrows in **B-D, F**. All flow cytometry data representative of at least 2 independent experiments.

**Supplementary Figure 6.**
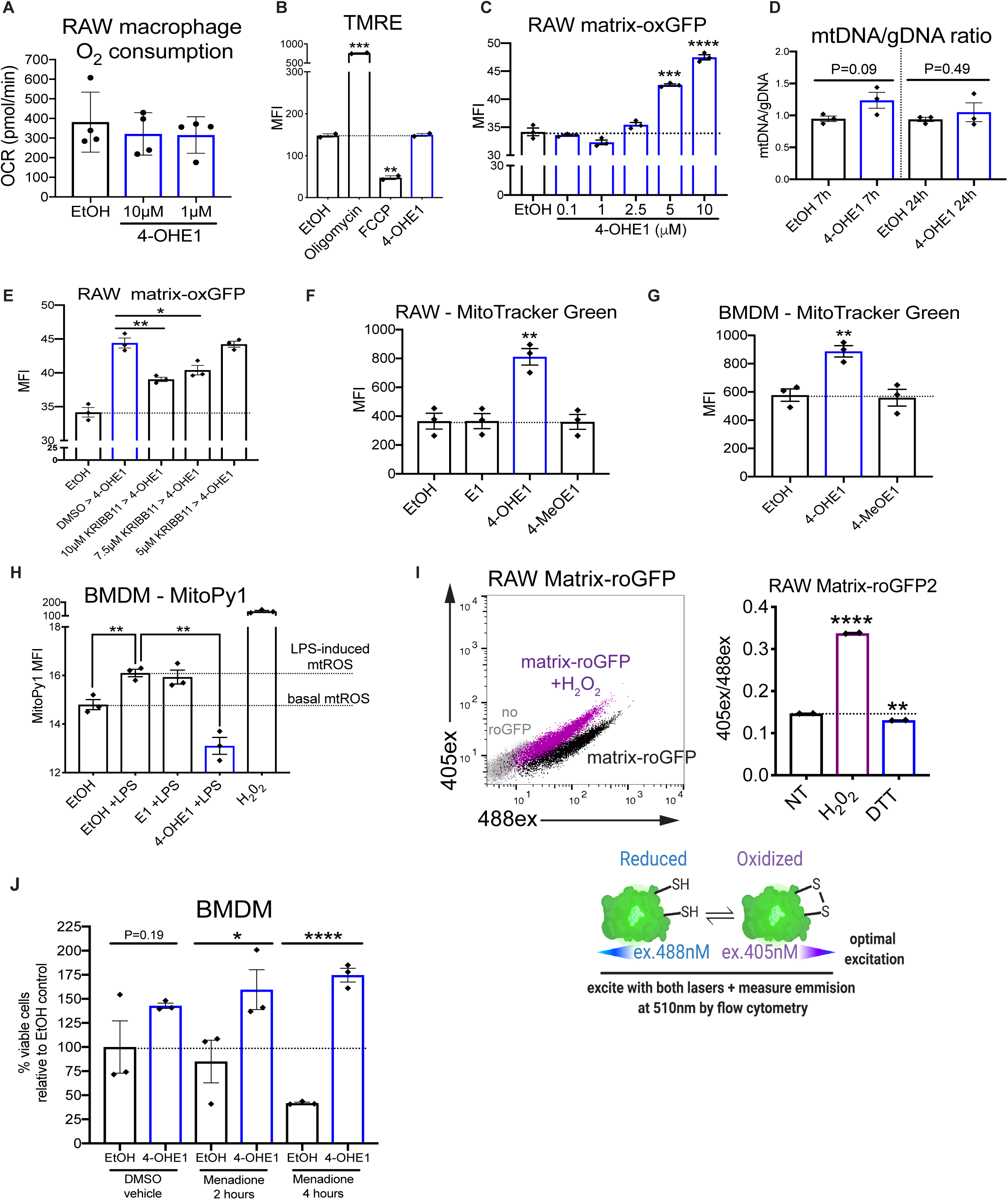
**A**. Seahorse respirometry measurement of oxygen consumption in RAW macrophages treated with EtOH vehicle control or indicated concentrations of 4-OHE1. Cells were treated immediately before loading into Seahorse XF24 analyzer, after which 5 repeated measurements per condition (with n=4 independent biological replicates per condition) were taken over 45 minutes and averaged.**B**. TMRE measurement of mitochondrial membrane potential in RAW macrophages treated with EtOH vehicle control, Oligomycin (5μM), FCCP (5μM), or 4-OHE1 (5μM) for 20 minutes before TMRE staining and flow cytometry.**C**. RAW matrix-oxGFP macrophages were treated with EtOH vehicle control or indicated concentrations of 4-OHE1 for 8 hours and matrix-oxGFP fluorescence quantified by flow cytometry.**D**. Mitochondrial DNA/genomic DNA (mtDNA/gDNA) ratio in RAW macrophages treated with EtOH vehicle control or 5μM 4-OHE1 for indicated times (n=3 independent biological replicates per condition). **E**. RAW matrix-oxGFP macrophages were pretreated with DMSO vehicle control or indicated concentrations of KRIBB11 for 1 hour before treatment with 5μM 4-OHE1 for 8 hours. Matrix-oxGFP fluorescence was then quantified by flow cytometry. **FG**. RAW macrophages (**F**) and BMDMs (**G**) were treated with EtOH vehicle control or estrogens (5μM) for 7 hours before MitoTracker Green staining and flow cytometry. **H**.BMDMs pretreated for 1 hour with EtOH vehicle control or estrogens (5μM) before LPS stimulation (100ng/mL) for 6 hours and mitochondrial H2O2 measurement with MitoPY1 staining and flow cytometry. **I. left** – RAW matrix-roGFP emission at 510nM with 405nm UV laser (y-axis) versus 488nM blue laser (x-axis) excitation. Shift of untreated matrix-roGFP cells (black population) following H2O2 treatment (1mM, purple population) for 10 minutes demonstrates ability to detect matrix-roGFP oxidation with flow cytometry. **right** – RAW matrix-roGFP 405nm laser excitation/488nM laser excitation ratio (405ex/488ex) after 10 minutes of H2O2 (1mM) and DTT (10mM) treatment.**F**. BMDMs treated overnight with EtOH or 5μM 4-OHE1 and treated the following day with DMSO vehicle control (4 hours) or 50μM menadione for (2,4 hours) before viability was assessed by flow cytometry. All bar graph data represented as mean ± SEM. For flow cytometry, n=2 or 3 independent biological replicates per condition. *P<0.05, **P<0.01, ***P<0.001,****P<0.0001 by unpaired, two-sided Student’s T Test (planned comparisons) except for **F** and **G** (one-way ANOVA with Dunnett’s test versus EtOH control sample).

Two hallmarks of mitohormesis are mitochondrial biogenesis, and increased mitochondrial chaperone activity^17, 18^. To determine if hydroxyestrogen-driven mitochondrial stress triggered these mitohormesis hallmarks, we expressed a mitochondrial matrix-targeted, oxidation-resistant green fluorescent protein^80^ in RAW macrophages, creating the RAW matrix-oxGFP reporter cell line. Treatment with 4-OHE1, but not E1 or 4-MeOE1, induced a steady increase in RAW matrix-oxGFP fluorescence at 8 and 24 hours as measured by flow cytometry (**Fig.6A**), an effect that was dose-dependent (**Supplementary Fig.6C**). Quantification of the mitochondrial DNA/genomic DNA (mtDNA/gDNA) ratio in RAW macrophages treated with 4-OHE1 revealed no significant increase (**Supplementary Fig.6D**). As matrix-oxGFP is a mitochondrial chaperone client that needs to be folded upon mitochondrial import^81^, this suggests RAW matrix-oxGFP cells are primarily reporting mitochondrial chaperone activity. Pharmacologically blocking HSF1 transcriptional activity with KRIBB11^82^ blunted increased matrix ox-GFP fluorescence in response to 4-OHE1, suggesting HSF1-dependent chaperone expression contributes to this mitohormetic response^18, 41^ (**Supplementary Fig.6E**). We do note that this increase in mitochondrial chaperone activity occurs concurrent with an increase in mitochondrial volume, as 4-OHE1, but not E1 or 4-MeOE1, increased MitoTracker Green signal in both RAW macrophages and BMDMs (**Supplementary Fig.6F,G**). Together, these results suggest hydroxyestrogen-driven mitochondrial stress triggers an adaptive increase in mitochondrial chaperone activity to protect and enhance mitochondrial proteostasis.

Another hallmark of mitohormesis is mitochondrial oxidative stress resistance (OSR)^17, 18, 83^; that is, a transient increase in mitochondrial stress triggers adaptations that both lower steady-state levels of mtROS at later timepoints, and provide defense against subsequent oxidative stress challenge. Like all cells, macrophages produce mtROS from the mitochondrial electron transport chain (ETC), and LPS stimulation enhances mtROS production for bactericidal purposes^8^. Thus, we investigated how hydroxyestrogen-driven mitochondrial stress affected mtROS levels in macrophages at later timepoints using the mitochondrial-targeted H2O2 sensor MitoPy1^84^. RAW macrophages treated for 7 hours with 4-OHE1, but not E1 or 4-MeOE1, showed reduced MitoPy1 fluorescence by flow cytometry, suggesting decreased basal mtROS levels (**Fig.6B**, left). In control one-hour EtOH pretreated cells, 6 hour LPS stimulation enhanced mtROS levels as expected (**Fig.6B**, right). However, in cells pretreated for one hour with hydroxyestrogens, but not with their precursor or methylated metabolites, there was a significant decrease in LPS-induced mtROS levels. This specific reduction in LPS-induced mtROS by hydroxyestrogen pretreatment also occurred in BMDMs (**Supplementary Fig.6H**).

To corroborate these effects, we expressed and targeted redox sensitive-GFPs(roGFPs)^85^ to the cytosol (cyto-roGFP), mitochondrial inner membrane space (IMS-roGFP), and mitochondrial matrix (matrix-roGFP) to quantify subcellular redox status in RAW macrophages (**Fig.6C**, left). Treatment of these cells with H2O2 and DTT confirmed our ability to detect changes in the redox state of roGFPs by flow cytometry (**Supplementary Fig.6I**). LPS stimulation for 6 hours revealed oxidation of IMS-roGFP, but not matrix-roGFP or cyto-roGFP, the latter of which showed a shift to a reduced state (**Fig.6C**, right). This demonstrates the ability of the roGFPs to monitor compartment-specific redox changes in LPS-stimulated macrophages, and supports the hypothesis that LPS-induced mtROS is primarily produced by Complex III into the IMS^12, 86, 87^. In agreement with our MitoPY1 data, pretreatment with hydroxyestrogens, but not their precursor or methylated metabolites, significantly reduced IMS-roGFP oxidation by LPS (**Fig.6D**). Together, these results suggest hydroxyestrogen-driven mitochondrial stress triggers mitohormetic mitochondrial OSR in macrophages, reducing endogenous mtROS levels.

We also tested whether this mitohormetic mitochondrial OSR could provide macrophages defense against normally toxic levels of pharmacologically-generated mtROS (**Fig.6E**, left schematic). Macrophages were treated overnight with EtOH or 4-OHE1, and viability was assessed the next day by flow cytometry. The majority of the EtOH and 4-OHE1-treated macrophages fell into a live forward scatter/side scatter (FS/SS) gate containing viable cells as assessed by DAPI exclusion (**Fig.6E**, right, top row). We then treated these macrophages with DMSO or a high dose of menadione, a redox cycling quinone that produces mitochondrial superoxide^83^. After 2 hours, an apoptotic cell population appeared in our EtOH control cultures, indicated by their FS/SS shift and inability to exclude DAPI; however, this population did not appear in our 4-OHE1-treated macrophages (**Fig.6E**, right, bottom row). We confirmed this resistance to menadione toxicity was specifically conferred by 4-OHE1, and not E1 or 4-MeOE1 (**Fig.6F)**, and that it occurred in BMDMs (**Supplementary Fig.6J**). Thus, mitohormesis in response to hydroxyestrogen-driven mitochondrial stress confers macrophages increased mitochondrial OSR and lasting “vaccine-like” protection^18^ against subsequent oxidative stress challenge with toxic levels of pharmacologically-generated mtROS.

### LPS-driven mitochondrial stress triggers mitohormesis identical to that observed in hydroxyestrogen-treated macrophages

Having characterized the mitohormetic response to hydroxyestrogen-driven mitochondrial stress, we next wondered what this pharmacologically-induced mitohormesis could teach us about how macrophages respond to LPS-driven mitochondrial stress. Oxidative damage, GST depletion, and upregulation of cytoprotective genes occurs rapidly (1-6 hours) following LPS treatment^12, 13^, likely in response to a combination of increased mitochondrial oxygen consumption and mtROS production^8, 12, 56, 57^, and increased mitochondrial production of electrophilic itaconate^88,89^. However, whether LPS-driven oxidative and electrophilic mitochondrial stress triggers mitohormesis is unknown.

To test the similarity between the mitochondrial stress caused by 4-OHE1 and LPS at the transcriptional level, we performed RNA-seq on BMDMs treated for 6 and 24 hours with either 4-OHE1 or LPS. This revealed an extremely high degree of overlap for both activated and repressed genes at each timepoint (**Fig.7A, Supplementary Table 4**). Chi-square tests confirmed such overlap would be extremely unlikely by chance (**Supplementary Table 5)**, suggesting 4-OHE1 very closely mimics physiological oxidative and electrophilic mitochondrial stress signals induced by LPS. GO analysis of genes upregulated by both 4-OHE1 and LPS at 6 and 24 hours revealed enrichment for categories including “Regulation of cellular response to stress”, “Protein folding”, and “Detoxification of ROS” (**Fig.7B, Supplementary Fig.7A)**. Examination of genes in these categories revealed concerted upregulation of HSF1-regulated molecular chaperones (HSPs = Heat Shock Proteins) that control mitochondrial protein folding, and enzymes of the peroxiredoxin/thioredoxin system that scavenge mitochondrial H2O2, by both 4-OHE1 and LPS (**Fig.7C, Supplementary Fig.7B**). 4-OHE1/LPS co-treatment further enhanced upregulation of many of these genes (**Fig.7D)**.

**Figure 7.**
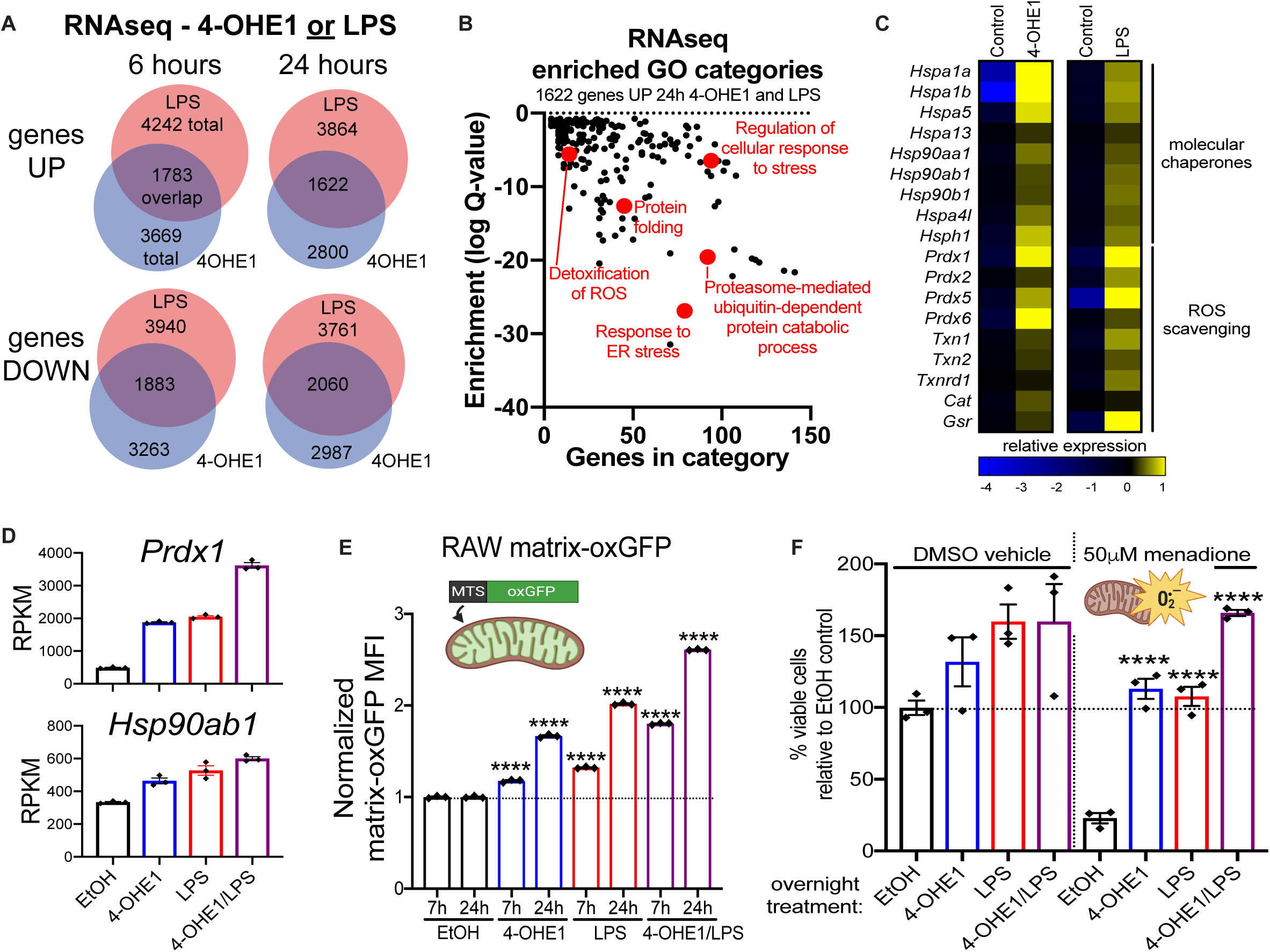
LPS-driven mitochondrial stress triggers mitohormesis identical to that observed in hydroxyestrogen-treated macrophages. **A**. BMDMs were treated with 4-OHE1 (5μM) or LPS alone for 6 or 24h, and RNA-seq was performed. Venn diagrams show overlap between genes significantly upregulated or downregulated by either treatment relative to controls. **B**.GO analysis of 1622 genes upregulated by both 4-OHE1 and LPS at 24h. **C**. Heatmap showing relative expression* of select genes upregulated by both 4-OHE1 and LPS at 24h. (*DESeq2 counts centered on the mean of each gene) **D**.Expression of *Prdx1* and *Hsp90ab1* (RPKM) at 24h from RNA-seq. **E**.Matrix-oxGFP RAW macrophages treated with EtOH, 4-OHE1 (5μM), LPS, or both, for indicated times and matrix-oxGFP florescence quantified by flow cytometry.****P<0.0001 by one-way ANOVA with Dunnett’s test versus time-matched EtOH control sample. **F**.Menadione resistance assay in BMDMs treated overnight (18-24h) with EtOH, 4-OHE1 (5μM), LPS, or both, before treatment with DMSO control or menadione (50μM, 4h) and viability assessment by flow cytometry. ****P<0.001 by one-way ANOVA with Dunnett’s test versus EtOH +Menadione sample. RNA-seq RPKM and flow cytometry data represented as mean ± SEM. Each data point is an independent biological replicate (n=3 for each condition). Flow cytometry data representative of 2 independent experiments.

**Supplementary Figure 7.**
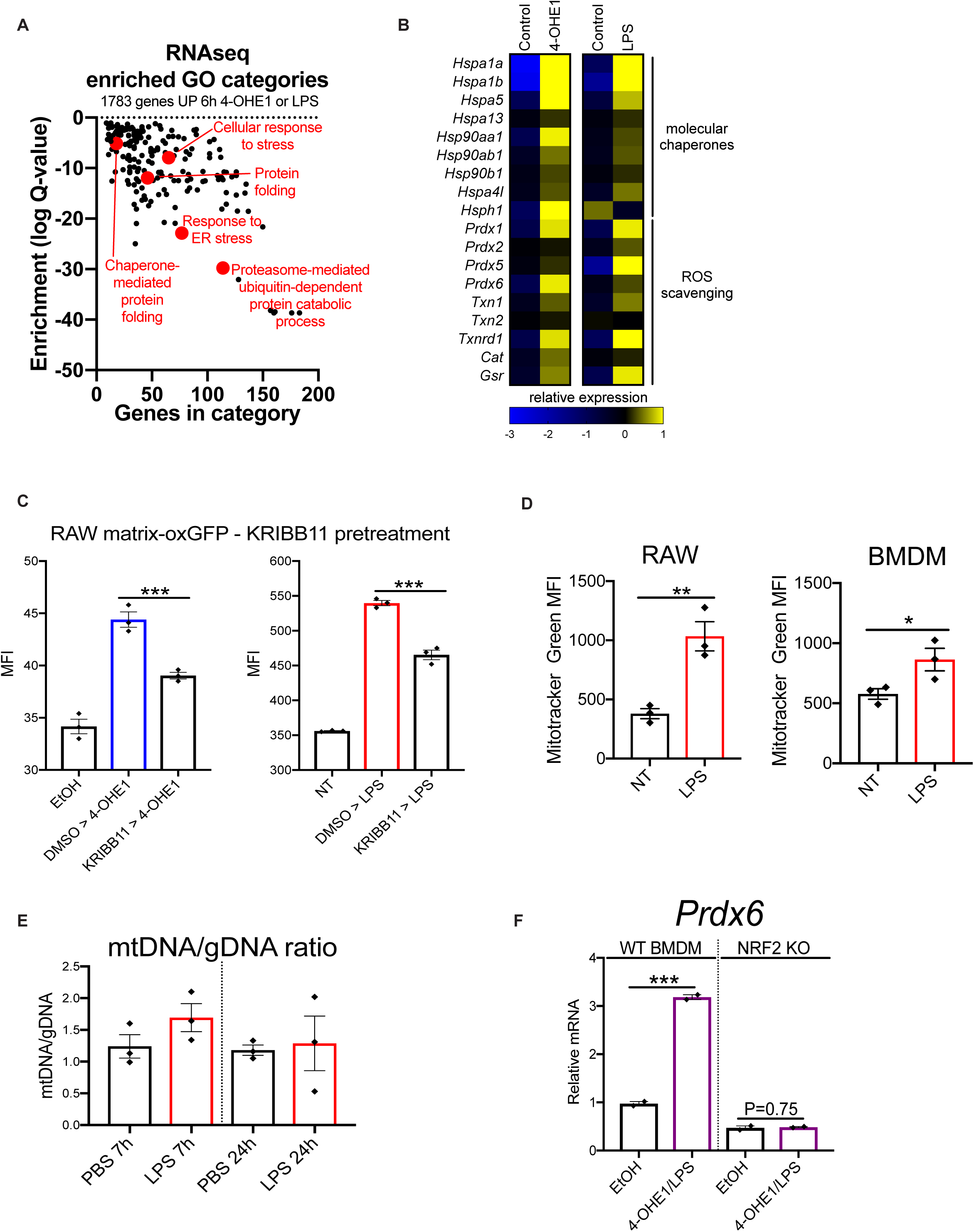
**A**.GO analysis of 1622 genes upregulated by both 4-OHE1 and LPS at 6 hours in BMDMs. **B**.Heatmap showing relative expression* of select genes upregulated by both 4-OHE1 and LPS at 6 hours in BMDMs. (*DESeq2 counts centered on the mean of each gene) **C**.RAW matrix-oxGFP macrophages pretreated with DMSO vehicle control or 10μM KRIBB11 for 1 hour before treatment with 4-OHE1 (5μM, left) or LPS (100ng/mL, right) for 8 hours. Matrix-oxGFP fluorescence was then quantified by flow cytometry.***P<0.001 by unpaired, two-sided Student’s T Test (planned comparisons). n=3 independent biological replicates per condition. **D**.MitoTracker Green signal in RAW macrophages and BMDMs measured by flow cytometry after 24h LPS simulation. **P<0.01 **P<0.01 by unpaired, two-sided Student’s T Test. n=3 independent biological replicates per condition. **E**.Mitochondrial DNA/genomic DNA (mtDNA/gDNA) ratio in RAW macrophages treated with PBS vehicle control or LPS for indicated times (n=3 independent biological replicates per condition). **G**. WT (left) and Nrf2 KO (right) BMDMs were treated with EtOH vehicle control, or 4-OHE1/LPS (5μM/100ng/mL) for 7 hours before *Prdx6* qPCR. ***P<0.001 by unpaired, two-sided Student’s T Test. n=2 independent biological replicates per condition. All bar graph data represented as mean ± SEM.

Given the highly similar transcriptional response to mitochondrial stress driven by 4-OHE1 and LPS, we next tested if, like 4-OHE1, LPS-driven mitochondrial stress triggered mitohormetic adaptations in macrophages. With regards to mitochondrial chaperone activity, treatment of RAW matrix-oxGFP reporter cells with LPS drove a progressive increase in fluorescence in a manner identical to 4-OHE1 treatment(**Fig.7E**). And like stress-induced gene expression, 4-OHE1/LPS co-treatment enhanced this effect. This suggests that the increased expression of molecular chaperones in response to both 4-OHE1 and LPS identified in our transcriptional profiling contributes to increased mitochondrial chaperone activity, a hallmark of mitohormesis. HSF1 transcriptional inhibition with KRIBB11 blunted increased matrix-oxGFP fluorescence in response to both 4-OHE1 and LPS, consistent with HSF1 regulating chaperone expression (**Supplementary Fig.7C**). And like 4-OHE1, LPS enhanced MitoTracker Green signal in macrophages without a significant increase in mtDNA content (**Supplementary Fig.7D and 7E**), suggesting an increase in mitochondrial volume.

To determine if, like 4-OHE1, LPS-driven mitochondrial stress triggered mitohormetic mitochondrial OSR, we repeated our menadione toxicity resistance experiments. 24 hour treatment with either 4-OHE1 or LPS increased BMDM viability relative to EtOH control cultures, with 4-OHE1/LPS co-treatment enhancing this effect (**Fig.7F**, left). Treatment of EtOH control BMDMs with menadione significantly decreased viability; however, both 4-OHE1- and LPS-treated macrophages were resistant to menadione-induced toxicity, with 4-OHE1/LPS co-treatment enhancing resistance (**Fig7F**, right). This suggests the increased expression of mtROS scavenging enzymes in response to both 4-OHE1 and LPS identified in our transcriptional profiling contributes to mitohormetic mitochondrial OSR. Induction of mtROS scavengers including *Prdx6* by 4-OHE1/LPS was impaired in NRF2 KO BMDMs, suggesting a role for NRF2 in mitohormetic OSR (**Supplementary Fig.7F**). Taken together, these data demonstrate that 4-OHE1-driven mitochondrial stress closely mimics physiologically relevant LPS-driven mitochondrial stress, and that both trigger classic mitohormetic adaptations in macrophages.

### Mitohormesis in macrophages includes metabolic reprogramming that enforces an LPS-tolerant state

Following primary LPS exposure, macrophages transition to an LPS-tolerant state where proinflammatory gene induction is refractory to upregulation by secondary LPS treatment (**Fig.8A**). Many mechanistic explanations for LPS tolerance have been proposed^90, 91^, including upregulation of negative regulators of TLR4 signaling, and production of anti-inflammatory cytokines including IL-10. However, recent evidence suggests that suppression of mitochondrial oxidative metabolism following LPS exposure limits acetyl-CoA production required for proinflammatory gene transcription^57^ (**Fig.8A**). ETC inhibition by LPS-induced nitric oxide (NO) production has been proposed to drive this suppression^92^; however, macrophages unable to produce NO still show significant suppression of mitochondrial oxidative metabolism following LPS treatment^93^. Metabolic reprogramming is often a part of mitohormetic responses to mitochondrial stress, as a shift away from mitochondrial oxidative metabolism towards aerobic glycolysis provides a damaged mitochondrial network an opportunity to recover from stress, while simultaneously augmenting ATP and NADPH production for energy and antioxidant defense, respectively^16, 17, 94-99^. Thus, we wondered if the mitochondrial stress-induced mitohormesis we observed in LPS-treated macrophages, which includes increased mitochondrial chaperone activity and OSR, also includes metabolic reprogramming that enforces tolerance via suppression of mitochondrial oxidative metabolism (**Fig.8A**). In other words, we hypothesized that mitochondrial stress is a key signal that triggers the transition from an LPS-responsive to LPS-tolerant state via mitohormetic metabolic reprogramming.

**Figure 8.**
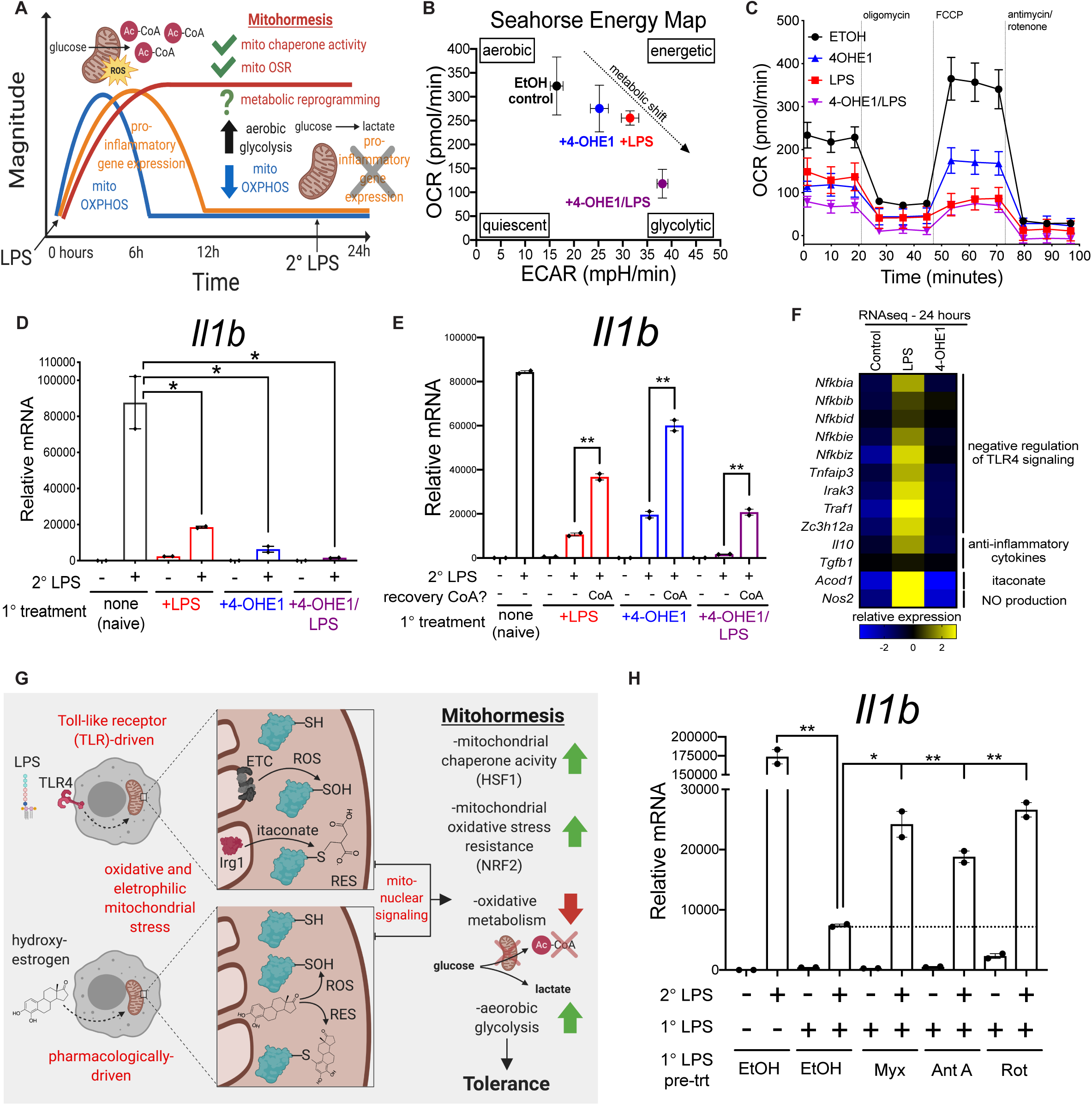
Mitohormesis in macrophages involves metabolic reprogramming that enforces an LPS-tolerant state. **A**.Schematic describing how mitochondrial oxidative metabolism (blue line) supports proinflammatory gene expression (orange line) after LPS treatment, but is suppressed as macrophages transition to an LPS-tolerant state where proinflammatory genes are refractory to upregulation by secondary LPS exposure. Mitohormetic adaptations (red line) occur in parallel with this process, but whether suppression of mitochondrial oxidative metabolism is a coincident mitohormetic adaptation is unknown. **B**.Seahorse energy map plotting basal oxygen consumption rate (OCR) versus basal extracellular acidification rate (ECAR) in RAW macrophages treated overnight (18-24h) with EtOH, 4-OHE1 (5μM), LPS, or both. **C**.Seahorse mitochondrial stress test in RAW macrophages treated overnight (18-24h) with EtOH, 4-OHE1 (5μM), LPS, or both. **D**.*Il1b* qPCR in RAW macrophages treated overnight (18-24h) with EtOH, 4-OHE1 (5μM), LPS, or both before treatments were washed out and cells allowed to recover (1-2h) before secondary LPS stimulation for 6h. **E**.Same as **D**, with CoA (2.5mM) provided to cells during the washout/recovery period before secondary LPS stim. **F**.Relative expression* of select genes in BMDMs treated with EtOH, 4-OHE1 (5μM), or LPS for 24h (*log2-transformed RPKM centered on the mean of each gene). **G**. Model showing how oxidative and electrophilic stress in LPS-stimulated macrophages drives mitohormesis and tolerance, and how hydroxyestrogens co-opt this stress response. **H**. RAW macrophages were pretreated 1h with vehicle or ETC inhibitors, then treated overnight (18-24h) with primary LPS before treatments were washed out and cells allowed to recover (1-2h) before secondary LPS stimulation for 6h and *Il1b* qPCR. All qPCR data represented as mean ± SEM. Each data point is an independent biological replicate (n=2 for each condition). *P<0.05, **P<0.01 by unpaired, two-sided Student’s T Test versus the indicated condition (planned comparisons). All qPCR data representative of 2 independent experiments except for CoA rescue, which needs to be repeated. Seahorse data representative of 2 independent experiments with n=5 independent biological replicates per condition. Seahorse data represented as mean ± SEM. Energy map represents the average of 3 repeated basal OCR measurements.

**Supplementary Figure 8.**
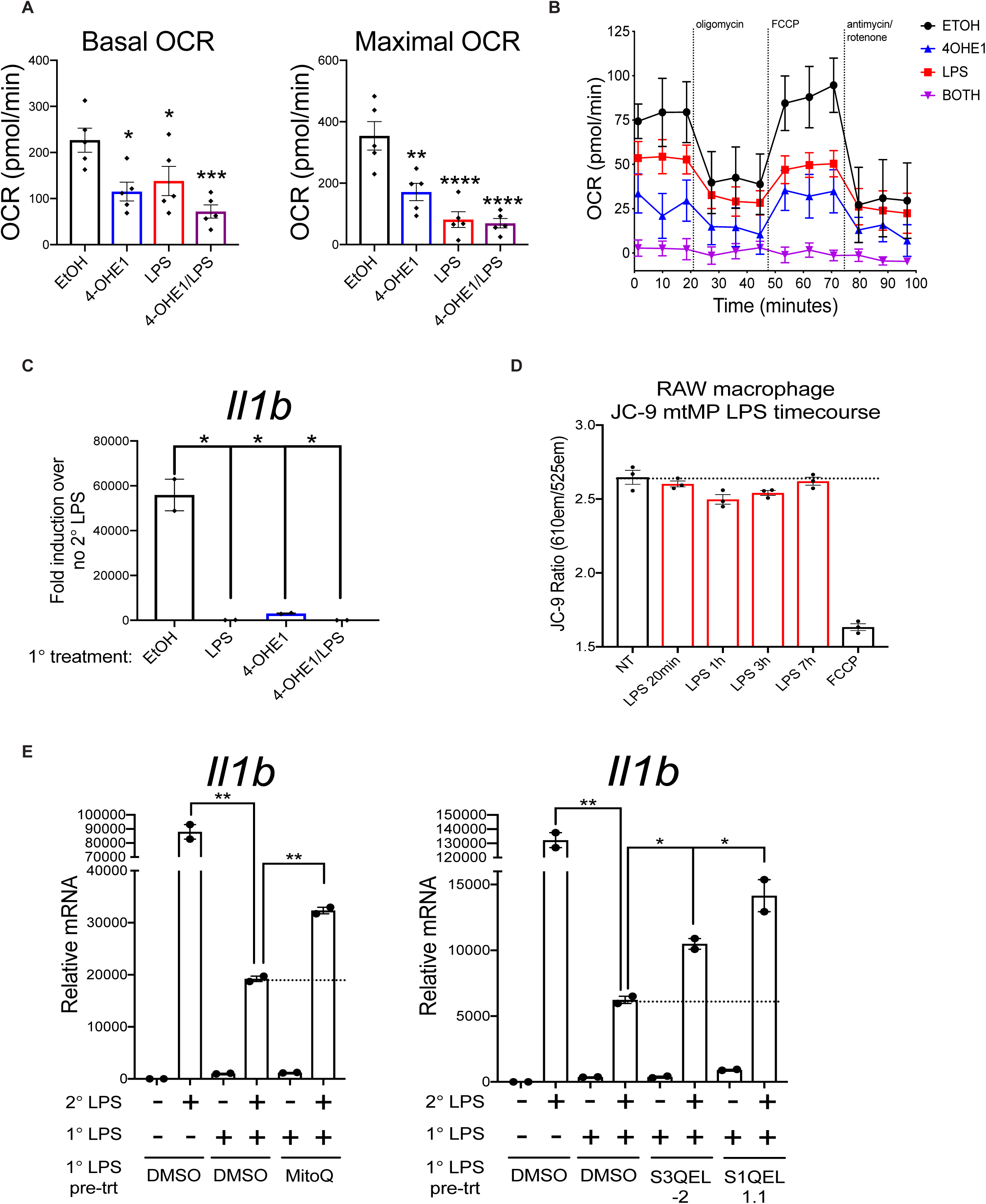
**A**. Basal and maximal OCR from RAW macrophage Seahorse Mito Stress Test in **Fig.8c**. Data is average of 3 repeated basal OCR measurements taken prior to oligomycin injection, or following FCCP injection (maximal), respectively. n=5 independent biological replicates per condition. *P<0.05, **P<0.01, ***P<0.001, ****P<0.0001 by one-way ANOVA with Dunnett’s test versus EtOH control sample.**B**. Seahorse mitochondrial stress test in BMDMs treated overnight (18-24 hours) with EtOH, 4-OHE1 (5μM), LPS (100ng/mL), or both. Data represented as mean ± SEM. n=5 independent biological replicates per condition.**C**. *Il1b* qPCR in BMDMs treated overnight (18-24 hours) with EtOH, 4-OHE1 (5μM), LPS (100ng/mL), or both, before treatments were washed out and cells allowed to recover (1-2 hours) before secondary LPS stimulation (100ng/mL) for 6 hours. Data is plotted as +/- LPS fold induction for cells with the same primary overnight treatment. n=2 independent biological replicates per condition. *P<0.05 by unpaired, two-sided Student’s T Test versus EtOH control (planned comparison).**D**. Flow cytometry measurement of mitochondrial membrane potential (mtMP) in RAW macrophages treated with LPS for indicated times and stained with JC-9.**E**. RAW macrophages were pretreated 1h with vehicle or inhibitors of mtROS production, then treated overnight (18-24h) with primary LPS before treatments were washed out and cells allowed to recover (1-2h) before secondary LPS stimulation for 6h and *Il1b* qPCR. *P<0.05, **P<0.01 by unpaired, two-sided Student’s T Test. All bar graph data represented as mean ± SEM. n=2 independent biological replicates for qPCR data.

If this hypothesis is true, then mitochondrial stress alone, separate from other TLR4-dependent events, should be sufficient to trigger metabolic reprogramming to an LPS-tolerant state via mitohormesis. Given how closely 4-OHE1-driven mitochondrial stress mimicked physiological LPS-driven mitochondrial stress, we tested if 4-OHE1 was sufficient to trigger mitohormetic metabolic reprogramming away from mitochondrial oxidative metabolism and towards aerobic glycolysis. RAW macrophages were treated with EtOH vehicle control, 4-OHE1, LPS, or both 4-OHE1/LPS for a tolerizing duration (18-24h) before treatments were washed out and cells subjected to Seahorse respirometry. Plotting basal extracellular acidification rate (ECAR, a proxy for glycolysis) versus basal oxygen consumption rate (OCR, a proxy for mitochondrial oxidative metabolism) to create an energy map showed that while naïve EtOH control macrophages are relatively aerobic, both 4-OHE1 and LPS treatment shifted the macrophages away from mitochondrial metabolism and towards aerobic glycolysis, with 4-OHE1/LPS cotreatment inducing an even stronger metabolic shift (**Fig.8B**). Seahorse mitochondrial stress testing revealed treatment with either 4-OHE1 or LPS alone was sufficient to significantly reduce basal and maximal OCR, with 4-OHE1/LPS cotreatment causing an even stronger reduction (**Fig.8C, Supplementary Fig.8A**). A similar response occurred in BMDMs (**Supplementary Fig.8B**). Thus, 4-OHE1-driven mitochondrial stress is sufficient to trigger mitohormetic metabolic reprogramming in a manner essentially identical to that observed in LPS-treated macrophages.

To test if the 4-OHE1 metabolically reprogrammed macrophages were LPS-tolerant, RAW macrophages were treated with EtOH, 4-OHE1, LPS, or both 4-OHE1/LPS for a tolerizing duration before treatments were washed out and cells allowed to recover. These macrophages were then either left untreated, or LPS-stimulated for 6 hours followed by *Il1b* qPCR (**Fig.8D**). While naïve macrophages responded robustly to LPS, cells treated overnight with primary LPS showed classic tolerance and impaired *Il1b* upregulation in response to secondary LPS. Similarly, cells treated overnight with 4-OHE1 also displayed impaired *Il1b* induction, demonstrating that 4-OHE1-induced metabolic reprogramming coincides with transition to an LPS-tolerant state. Finally, in agreement with 4-OHE1/LPS cotreatment driving a stronger metabolic shift than either treatment alone, macrophages co-treated with 4-OHE1/LPS overnight displayed an even more severely impaired secondary LPS response. CoA supplementation during the washout and recovery period boosted LPS responsiveness in both LPS- and 4-OHE1-tolerized RAW macrophages, suggesting impaired CoA/ acetyl-CoA homeostasis is a common feature of these tolerized states (**Fig.8E**). 4-OHE1 also induced tolerance in BMDMs (**Supplementary Fig.8C**). Together, these data demonstrate that in addition to increased mitochondrial chaperone activity and OSR, 4-OHE1-induced mitohormesis involves metabolic reprogramming and transition to an LPS-tolerant state essentially identical to that induced by LPS. Our RNA-seq data revealed that unlike LPS, 4-OHE1-induced LPS tolerance occurs in the absence of transcriptional upregulation of negative regulators of TLR4 signaling such as *Tnfaip3, Il10*, and without *Nos2* induction (**Fig.8F**). Thus, mitohormetic metabolic reprogramming to an LPS-tolerant state can be uncoupled from TLR4 signaling, and mitochondrial stress is a sufficient signal able to trigger this reprogramming in the absence of other TLR4-dependent events that have been proposed to enforce tolerance.

We propose a model where mtROS- and/or mtRES-induced mitochondrial stress drive mitohormesis, and in turn, a state of “mitohormetic tolerance” (**Fig.8G**). One prediction of this model is that reducing mtROS during primary LPS exposure would impair the oxidative stress signaling that leads to tolerance, and enhance secondary LPS responsiveness. mtMP has been reported to increase following LPS treatment, inducing reverse electron transport (RET) to drive mtROS production^75^; however, this TMRE-based measurement, made at 24 hours post-LPS treatment, was not normalized to mitochondrial volume, which increases post-LPS treatment (**Supplementary Fig.7D**). Ratiometric mtMP measurements with JC-9 revealed no increase immediately post-LPS treatment (**Supplementary Fig.8D**), suggesting LPS-induced mtROS is a product of increased ETC forward flux and mitochondrial oxygen consumption that has been measured in the 0.5-2 hours following LPS treatment^12, 57^. Inhibiting this ETC flux with rotenone, antimycin A, or myxothiazol during primary LPS exposure resulted in a partial, but significant, rescue of secondary LPS responsiveness (**Fig.8H**). A similar effect was observed when MitoQ, S1QEL1.1, and S3QEL-2, molecules that reduce mtROS without inhibiting the ETC, were present during primary LPS exposure (**Supplementary Fig.8E**). These data suggest mtROS is partially responsible for mitohormetic tolerance and restraining macrophage inflammatory responsiveness.

## Discussion

Whether intrinsically-generated oxidative and electrophilic mitochondrial stress affects macrophage function is unknown. Moreover, the mechanisms controlling macrophage tolerance are unclear. We identified hydroxyestrogens as potent anti-inflammatories, and through study of their pharmacological effects, we demonstrate that both hydroxyestrogen- and LPS-driven mitochondrial stress trigger a set of stress adaptations known as mitohormesis. One of these adaptations, the suppression of mitochondrial oxidative metabolism, suppresses the macrophage response to secondary LPS, revealing mitohormesis as a stress response that enforces of LPS tolerance to prevent excessive inflammation. Thus, in addition to their anti-microbial roles, mtROS and itaconate are signaling molecules that induce a negative feedback loop to restrain inflammation via mitohormesis, a process that can be pharmacologically targeted (**Fig.8G**).

Whether the acute depletion of CoA and acetyl-CoA by hydroxyestrogens occurs through targeted inhibition of a specific mitochondrial protein, or through more general oxidative and electrophilic mitochondrial stress, will require further investigation. In support of the latter mechanism, we show the repression of LPS-induced *Il1b* in macrophages by multiple immunomodulatory electrophiles can be overcome by providing exogenous acetyl-CoA (**Fig.5I**). Interestingly, much lower doses of more lipophilic electrophiles are sufficient to repress *Il1b* to levels similar to 100μM DEM, suggesting the ability to cause stress locally in mitochondrial membranes dictates anti-inflammatory potency of these small molecules.

Regarding macrophage mitohormesis, while we provide evidence that HSF1 and NRF2 regulate specific adaptations (increased mitochondrial chaperone activity and OSR, respectively), the identity of the transcription factor(s) coordinating suppression of mitochondrial oxidative metabolism remains unclear. ATF4 is an attractive candidate, as its *C. elegans* orthologue ATFS-1 simultaneously suppresses mtETC genes and upregulates glycolytic genes in response to mitochondrial stress^95, 96^. For LPS-induced mitohormesis, the relative contributions of mtROS, itaconate, and other RES (e.g. 4-hydroxynonenal) in triggering mitohormesis and enforcing tolerance will require further investigation. While we provide evidence that mtROS is partially required for mitohormetic tolerance, *Irg1*-deficient macrophages treated with LPS are resistant to suppression of mitochondrial oxidative metabolism^100^, suggesting multiple reactive molecules contribute to triggering mitohormesis. Overall, we speculate that this metabolic reprogramming involves both transcriptional changes, along with post-transcriptional alterations in mitochondrial composition by the ubiquitin-proteasome system, which can tune mitochondrial composition and function in response to oxidative and metabolic stress^101^. Importantly, whether ROS/RES induce mitohormetic tolerance in clinical situations of immunosuppression where restoring myeloid inflammatory responses would be beneficial (e.g. late-stage sepsis, cancer) will be important to investigate. For example, a HIF-1 signature of increased glycolytic gene expression is shared between myeloid cells from patients with sepsis or cancer^102^. Moreover, buildup of the RES methylglyoxal was recently found to suppress oxidative metabolism and inflammatory function in cancer-promoting myeloid cells^103, 104^. We propose myeloid cells performing high levels of aerobic glycolysis (glucose to lactate) represent exhausted myeloid cells unable to convert glucose-derived carbon into acetyl-CoA to epigenetically activate inflammatory responses.

From an anti-inflammatory therapeutic perspective, these findings demonstrate that beyond acute depletion of acetyl-CoA, inducing mitohormetic tolerance represents a mechanism by which the reactive, lipophilic hydroxyestrogens achieve lasting immunosuppression. We propose triggering mitohormesis in macrophages by causing oxidative and electrophilic mitochondrial stress with lipophilic electrophiles represents a novel therapeutic strategy to combat inflammation that works by co-opting the physiological response to mitochondrial stress that occurs naturally after primary LPS exposure and during transition to an LPS-tolerant state. In other words, by promoting eustress^105^ (i.e. moderate mitochondrial stress), a benefit is achieved (i.e. reduced inflammation). This concept of a mitochondria-targeted, pro-oxidant therapy to repress inflammation via triggering mitohormetic metabolic reprogramming stands in direct opposition to the concept of using mitochondria-targeted anti-oxidants as anti-inflammatories, which has been employed by others in the macrophage immunometabolism field^12, 75, 86, 87^

## Acknowledgements

We thank the following people for their essential contributions to this manuscript: Hector Nolla and Alma Valeros for assistance with cell sorting and flow cytometry; the Dillin Lab for JC-9 reagent; Dr. Ching Fang Chang, Pete Zushin, and Garret Dempsey for reagents, assistance with Seahorse respirometry, and helpful discussion; Dr. Hong Sik Yoo and Dr. Joseph Napoli for assistance with steroid extraction; members of the UC Berkeley Functional Genomics Laboratory and Vincent J. Coates Genomic Sequencing Laboratory for assistance with next-generation sequencing; Claudia Rosso, Kevin Shohat, Jordan Uyeki, and Daniel Tan of the UCLA Metabolomics Center for LC-MS data processing and analysis; members of the Welch Lab for assistance with SpeedVac; members of the Saijo Lab and UC Berkeley immunology community for helpful feedback; Dr. Andrew Dillin and Dr. Samantha Lewis for critical reading of the manuscript; Dr. Laura Lau for emotional support and baked goods.

## Author contributions

K.S. and G.T. conceptualized project. G.T., K.M.T., A.T.I., J.t.H., D.K.N., A.S., V.M.W., and K.S. procured funding and resources. G.T., K.M.T., B.F., A.T.I., and J.t.H. designed methodology. G.T., K.M.T., B.F., J.W., J.W., S.Z., R.K., S.L., A.T.I., and J.t.H. performed experiments and analyzed data. G.T. curated data and wrote manuscript.K.S. and G.T. edited manuscript with input from all authors.

## Methods

### Animals

All experiments were approved by the UC Berkeley Animal Care and Use Committee (ACUC) (AUP-2017-02-9539-1 to K.S.) and performed under the supervision of the UC Berkeley Office of Laboratory and Animal Care (OLAC). Nrf2 KO mice (Stock No. 017009) were purchased from Jackson Labs.

### Cell culture

Bone marrow-derived macrophages (BMDMs) were prepared by flushing leg and hip bones from 6-12-week-old mice, filtering cells through a 40μm filter, RBC lysis, and plating in DMEM supplemented with 20ng/mL murine M-CSF (Shenandoah) in 15cm Petri dishes (non-TC treated). Media/M-CSF was changed on days 3 and 6. Cells were harvested by scraping and plated in appropriate TC-treated multiwell plates/dishes for experiments on days 7-10. For initial estrogen metabolite experiments, phenol red-free DMEM (Corning 17-205-CV) and 10% charcoal-stripped FBS (HyClone SH30068.03) were used to eliminate estrogenic effects of phenol red and serum estrogens. All experiments after verifying that hydroxyestrogen effects were ER-independent were performed in DMEM with phenol red (Corning 10-013-CV) supplemented with 10% regular FBS (HyClone SH30071.03). Both DMEM/FBS formulations were supplemented with Pen/Strep (Gibco). All BMDM data presented are with cells derived from female mice, though we confirmed that hydroxyestrogens are anti-inflammatory in BMDMs derived from male mice. RAW 264.7 macrophages and HEK293T cells (both ATCC) were also cultured in DMEM with 10% FBS and Pen/Strep. The latter were used to produce lentivirus with psPAX2 (Addgene 12260), pMD2.G (Addgene 12259), and various lentiviral constructs described hereafter.

### Chemicals

All estrogens were from Steraloids Inc. (see Supplementary Table 1). Pam3CSK4 (tlrl-pms) and ODN (tlr1-1826-1) were from InvivoGen. LPS (L3024), poly IC (P0913), diethyl maleate (D97703), sodium acetate (S5636), celastrol (C0869), myxothiazol (T5580-1MG), S1QEL1.1 (SML1948-5MG), and S3QEL-2 (SML1554-10MG) were from Sigma. Coenzyme A (CoA) and acetyl-CoA (Ac-CoA) from both Sigma (C4780, A2056) and Cayman (16147, 16160) were tested, and both behaved similarly in their rescue of proinflammatory gene expression in hydroxyestrogen-pretreated macrophages. All *in vitro* LPS stimulations performed using 100ng/mL dose. Pam3CSK4, pIC, and ODN were used at 100ng/mL, 25μg/mL, and 1μM, respectively. MitoQ (89950) was from Cayman.

### Quantitative real-time PCR (qPCR)

Cells in multiwell plates were directly lysed in Trizol (Invitrogen) and total RNA isolated using Direct-zol kit (Zymo). cDNA was prepared with Superscript III (Invitrogen) and diluted in H2O for qPCR using ROX low SYBR FAST 2X mastermix (KAPA/Roche) and QuantStudio6 qPCR machine (Thermo Fisher) in 96-well plate fast run mode. Data was collected and analyzed using QuantStudio Real Time PCR Software v1.3. Primers (IDT, Supplemental Table 6) were designed to span exon-exon junctions in Primer-BLAST^1^ and determined to have one product via melt curve analysis. All data presented is normalized to *Hprt* reference gene expression using the delta-delta Ct method.

### RNA-seq

For LPS-stimulated BMDM dataset with different pretreatments (introduced in Fig.1C), n=2 independent biological replicates were pooled to generate total RNA samples. For vWAT macrophage dataset (introduced in Fig.2C), SVF from 2 or 3 mice was pooled, and two independent sorts of macrophages were done to generate total RNA samples. For 4-OHE1- and LPS-treated BMDM dataset (introduced in Fig.7A), n=3 independent biological replicates were used to generate total RNA samples

For library preparation, total RNA was converted into sequencing libraries using mRNA HyperPrep kit (KAPA/Roche) and custom Illumina-compatible unique dual index (UDI) adaptors (IDT). Libraries were quantified with Illumina Library qPCR quantification kit (KAPA/Roche) and pooled for sequencing on HiSeq4000 (Illumina). Sequencing reads were aligned to mm9 or mm10 using STAR^2^, and reads counted (raw counts and RPKM) using HOMER^3^. EdgeR^4^ and DESeq2^5^ were used for differential expression analysis using raw read counts, and all differential expression gene lists were generated using the cutoff parameters indicated in Supplemental Tables 1-6. Hierarchical clustering was performed using Cluster^6^ and visualized with Java TreeView. Heatmaps were produced from normalized expression data in either Java TreeView or GraphPad Prism 8, collapsing independent biological replicates into one column representing the relative expression for each indicated treatment condition. Gene ontology (GO) analysis was performed by inputting gene lists into Metascape^7^. Promoter motif finding from gene lists was performed using HOMER.

### Western blotting

Cells/mitochondrial fractions were lysed in RIPA buffer plus protease inhibitor cocktail (Roche) for 20 min on ice, followed by centrifugation 10,000g x 10 min at 4°C. Supernatant was quantified with DC Protein Assay (BioRad), and 15-30μg of soluble protein mixed with 5x NuPAGE loading dye and 2x NuPAGE reducing reagent (Life Technologies) and heated 10 min at 70°C. Samples were loaded on NuPAGE 4-12% Bis-Tris gels (Life Technologies) against SeeBlue Plus2 prestained protein ladder (Thermo Fisher) and ran in NuPAGE MOPS buffer (Life Technologies). Size-separated proteins were transferred to PVDF membrane (GE Healthcare) by semidry transfer, and membrane blocked with SuperBlock (Thermo Fisher) for 1 hour. Membranes were probed with primary antibodies overnight at 4°C, followed by washes with 0.1% TBS-T, and fluorophore-conjugated secondary antibody probing at room temperature for 1 hour. After 0.1% TBS-T washes, membrane fluorescence was visualized using Licor Odyssey imaging system. Primary antibodies: Tubulin (CP06, Cal Biochem), pro-IL-1β (AF-401-NA, R&D Systems), NRF2 (MABE1799, EMD Millipore), vinculin (sc-73614, Santa Cruz), VDAC (ab154856, Abcam). Secondary antibodies: AlexaFluor 680 conjugates were from Invitrogen. IRDye 800CW conjugates were from Rockland. All antibodies were diluted in 0.1% TBS-T with 5% BSA and 0.02% sodium azide for probing.

### Flow cytometry and cell sorting

All sample analysis was done on LSR II, LSR Fortessa, or LSR Fortessa X20 analyzers (BD). All cell sorting was performed on Influx or Aria Fusion sorters (BD). Data was collected using BD FACSDiva software and analyzed using FlowJo 10.5.2 (Tree Star).

### Pro-IL-1β ICS for flow cytometry

RAW macrophages were treated, harvested, and processed with FIX & PERM Cell Fixation & Permeabilization Kit (Thermo Fisher) in 96-well round-bottom plate. Primary antibody: pro-IL-1β (AF-401-NA, R&D Systems) diluted 1:50 in Solution B. Secondary antibody: anti-goat AlexaFluor 647 (Invitrogen) diluted 1:500 in Solution B. All staining done in 50μL volume. All washes performed with PBS.

### Acute *in vivo* inflammation

Male C57BL/6J mice (Jackson Labs) mice were injected intraperitoneally (IP) with EtOH vehicle control or estrogens (10mg/kg). One hour later, mice received IP injection of PBS or LPS (2mg/kg). After 3 hours, submandibular bleeding was performed to collect blood for measurement of serum IL-1β levels by ELISA (Invitrogen 88-7013-22). At 4 hours, mice were sacrificed and splenocytes isolated by crushing spleen through 40μm filter, performing RBC lysis, and lysing total splenocyte cell pellet in Trizol for RNA extraction, cDNA preparation, and qPCR for proinflammatory genes.

### Chronic *in vivo* inflammation

Male C57BL/6N mice (Charles River) mice were placed on high-fat diet (Research Diets Inc, D12492, 60 kcal% fat) and given subcutaneous EtOH vehicle control or estrogen injections (10mg/kg) every 6 days in rear flank. After 30 days, mice were sacrificed and visceral white adipose tissue (vWAT) isolated and weighed. Stromal vascular fraction (SVF) was then prepared from vWAT. Tissue was minced with scissors and incubated in DMEM with 0.1% collagenase (Sigma C6885) and 5% BSA for 1 hour at room temperature with gentle shaking. Digested tissue was filtered through a 70μm filter, RBCs lysed, and SVF stained for analysis and cell sorting on ice in FACs buffer (PBS with 10mM HEPES and 5% BSA) using standard techniques. After incubation in Fc Block (BD Biosciences), cells were stained with anti-CD45-PerCP-Cy5.5 (clone 30-F11, BioLegend), anti-F4/80-PE (clone BM8, eBioscience), and anti-CD11b-APC (clone M1/70, eBioscience). After washes, cells were resuspended in FACs buffer with DAPI (Thermo Fisher D1306, reconstituted according to manufacturer instructions, used at 1:100,000) for exclusion of dead cells. F4/80+CD11b+ vWAT macrophages were sorted directly into Trizol-LS (Invitrogen) for RNA isolation. Total live SVF cell counts were also determined using hemocytometer and Trypan Blue exclusion so that macrophage cellularity could be calculated from flow cytometry percentages.

### Mitochondrial fractions for steroid extraction and LC-MS

RAW macrophages (15-30 million total) were treated with EtOH vehicle control or 5μM 4-OHE1 for 1 hour before 1x PBS wash and harvest by scraping. For whole cell steroid extraction, 1 million macrophages were pelleted (800g x 5 minutes, 4°C) in a 1.5mL Eppendorf tube, flash frozen, and stored at -80°C. From remaining cells, mitochondrial fractions were isolated using a previously described differential centrifugation method^8^ with modification. Briefly, cells were resuspended in 2mL cell isolation buffer (IBc) in a 15mL conical and disrupted using a Bioruptor sonicator (Diagenode) with metal probe adaptor and short one-second pulses on “HI” setting. Cell disruption was tracked by flow cytometry scatter, with 10-15 pulses getting cells from 70% to <10% “live scatter” gate-positive. Mitochondrial fractions were then isolated by differential centrifugation, flash frozen, and stored at -80°C. Fractions were checked for mitochondrial protein enrichment versus whole cell lysates by western blot.

For steroid extraction, whole cell and mitochondrial fraction pellets were thawed at room temperature and resuspended in 1 mL of acetonitrile by pipetting and vortexing. The resulting homogenate was stored for 30 min at −20°C and then centrifuged for 5 min at 12000 × *g* and at 4°C. The supernatant was transferred to a glass round-bottom tube and evaporated under a N2 stream. The residue was resuspended in 2mL of 0.2M sodium acetate buffer (pH 5.0) and extracted with 10mL of hexane. The mixture was centrifuged for 2 min at 1200 × *g*, and the upper hexane layer was transferred to a new glass tube and evaporated under nitrogen with gentle heating at 25–30°C in a water bath (Model N-EVAP 112, Organomation Associates). The residue was reconstituted in 50μL of MeOH with vortexing. As a positive control to confirm this method could extract 4-OHE1, we also performed an extraction from 20μL of cell culture media to which 2μL of 5mM 4-OHE1 was added and immediately flash frozen.

Extracts and 4-OHE1 standard were analyzed using a liquid chromatography system (LC; 1200 series, Agilent Technologies, Santa Clara, CA) that was equipped with a reversed-phase analytical column (length: 150 mm, inner diameter: 1.0 mm, particle size: 5 µm, Viva C18, Restek, Bellefonte, PA). The LC system was connected in line with an LTQ-Orbitrap-XL mass spectrometer that was equipped with an electrospray ionization (ESI) source and operated in the positive ion mode (Thermo Fisher Scientific, Waltham, MA). Mass spectrometry data acquisition and processing were performed using Xcalibur software (version 2.0.7, Thermo). Injection volumes were as follows: 2μL for 1mg/mL 4-OHE1 standard (Steraloids) in MeOH, 5μL for whole cell extracts, and 2μL for mitochondrial extracts. The LC-MS is located in the QB3/Chemistry Mass Spectrometry Facility, on the campus of the University of California, Berkeley.

### isoTOP-ABPP

IsoTOP-ABPP analysis was performed as previously described^9^. Briefly, proteomes (prepared from BMDMs treated with EtOH or 1μM 4-OHE1 for 1 hour) were labeled with IAyne (100 μM) for 1 hour at room temperature, and subsequently treated with 100 μM isotopically light (control) or heavy (treated) TEV-biotin and click chemistry was performed as previously described^10^. Proteins were precipitated, washed, resolubilized and insoluble components were precipitated. Soluble proteome was diluted and labeled proteins were bound to avidin-agarose beads while rotating overnight at 4°C. Bead-linked proteins were enriched, then resuspended, alkylated with iodoacetamide, then washed and resuspended with sequencing grade trypsin overnight. Non-bead-linked tryptic peptides were washed away, and the TEV-biotin tag was digested overnight in TEV buffer containing and Ac-TEV protease at 29°C. Liberated peptides were diluted in water, acidified and stored at -80°C.

Analysis was performed as previously described using Multidimensional Protein Identification Technology (MudPIT) with an Orbitrap Q Exactive Plus mass spectrometer (Thermo Fisher)^9^. Data was extracted in the form of MS1 and MS2 files using Raw Extractor 1.9.9.2 (Scripps Research Institute) and searched against the UniProt mouse database using ProLuCID search methodology in IP2 v.3 (Integrated Proteomics Applications, Inc)^11^. ProLUCID data was filtered through DTASelect to achieve a peptide false-positive rate below 1% and cysteine residues were searched with a static modification for carboxyaminomethylation (+57.02146 Da) and up to two differential modifications for the light or heavy TEV tags (+464.28596 or +470.29977 Da, respectively).

### Metabolomics

BMDMs were cultured in DMEM (Corning 17-207-CV) supplemented with One Shot dialyzed FBS (Thermo Fisher), 1mM sodium pyruvate, 4mM L-glutamine, and 25mM glucose (all from Gibco). Cells were seeded into 6-well plates at 500,000 cells per well, and the next day treated with EtOH vehicle control or 5μM 4-OHE1. After two hours, media was aspirated and cells washed 1x with unsupplemented DMEM. Media was then replaced with DMEM described above except with 25mM ^13^C6-glucose (Cambridge Isotope Laboratories, CLM-1396-PK). After 30 minutes, media was aspirated, cells washed 1x with 1mL PBS, and moved to dry ice for addition of 1mL ice cold 80% MeOH. Cells were incubated at -80°C for 15 minutes, after which cells were scraped on dry ice and moved to Safe Lock 1.5mL Eppendorf tube (cat.022363204). After centrifugation 20,000g x 10 minutes at 4°C, supernatant was transferred to a new tube and evaporated overnight using a SpeedVac with no heating. Dried metabolite extracts were stored at -80°C before shipping on dry ice. MeOH extraction pellet was saved for protein quantification to be used in metabolite normalization.

Dried metabolites were resuspended in 50% ACN:water and 1/10^th^ was loaded onto a Luna 3um NH2 100A (150 × 2.0 mm) column (Phenomenex). The chromatographic separation was performed on a Vanquish Flex (Thermo Scientific) with mobile phases A (5 mM NH4AcO pH 9.9) and B (ACN) and a flow rate of 200μL/min. A linear gradient from 15% A to 95% A over 18 min was followed by 9 min isocratic flow at 95% A and reequilibration to 15% A. Metabolites were detected with a Thermo Scientific Q Exactive mass spectrometer run with polarity switching (+3.5 kV/− 3.5 kV) in full scan mode with an m/z range of 65-975. TraceFinder 4.1 (Thermo Scientific) was used to quantify the targeted metabolites by area under the curve using expected retention time and accurate mass measurements (< 5 ppm). Values were normalized to sample protein concentration. Relative amounts of metabolites were calculated by summing up the values for all isotopologues of a given metabolite. Fraction contributional (FC) of ^13^C carbons to total carbon for each metabolite was calculated as previously described^12^. Data analysis was performed using in-house R scripts.

### ChIP-seq

Approximately 30 million total BMDMs (3 independent biological replicates of 10 million cells each) were treated with EtOH vehicle control or 1μM hydroxyestrogen for 1 hour, followed by PBS or LPS (100ng/mL) stimulation for 30 minutes. Cells were then washed 2x with PBS and fixed with 0.67mg/mL DSG (Thermo Fisher) in PBS for 30 minutes at room temperature with shaking, followed by addition of paraformaldehyde (Electron Microscopy Sciences) to 1% and an additional 15 minutes of room temperature fixation. Fixation was quenched by adding glycine to 125mM and shaking 10 minutes, after which fixed cells were scraped and washed 2x with PBS. Cell pellet was resuspended in 1.5mL ice cold ChIP RIPA buffer (20mM Tris HCl pH 8.0, 150mM NaCl, 2mM EDTA, 0.1% SDS, 1% Triton X-100) with protease inhibitors (Roche) and sonicated with metal probe adaptor using Bioruptor (Diagenode) for 60 minutes (continuous cycles of 20 seconds ON/40 seconds OFF, “medium” setting). Samples were spun 20 minutes at maximum speed at 4°C in a benchtop centrifuge to remove insoluble material, and supernatant containing soluble sheared chromatin transferred to new tube, saving 1% volume for input library preparation. Immunoprecipitation was performed overnight at 4°C with 2μg of primary antibody (p65: Santa Cruz sc-372, H3K27ac: Abcam ab4729), or corresponding species-appropriate IgG control antibody (GenScript), conjugated to Protein A Dynabeads (Invitrogen). The next day, beads were captured with magnet on ice and washed with ice cold wash buffer II (20mM Tris HCl pH 8.0, 150mM NaCl, 2mM EDTA, 1% Triton X-100, 0.5% NaDOC) 3 times, wash buffer III (10mM Tris HCl pH 8.0, 250mM LiCl, 2mM EDTA, 1% NP-40, 0.5% NaDOC) 3 times, and TE with 50mM NaCl two times, all supplemented with protease inhibitors. Beads with antibody-target-DNA complexes were then resuspended in 200μL elution buffer (50mM Tris HCl pH 8.0, 10mM EDTA, 1% SDS) for 30 minutes at 37°C to elute immunoprecipitated complexes. 200μL eluent was collected, 10μL of 5M NaCl added, and DNA-protein crosslinks reversed overnight at 65°C. Next day, samples were treated with RNAse A (Thermo Fisher) for 1 hour at 37°C, Proteinase K (NEB) for 1 hour at 50°C, and DNA fragments recovered with Zymo ChIP DNA Clean & Concentrator kit.

Sequencing libraries were prepared from both input chromatin, and chromatin recovered from immunoprecipitations, using in-house protocol. Briefly, DNA was blunted, A-tailed, and ligated to Illumina-compatible NEXTFlex sequencing adaptors (Bioo). Libraries were PCR amplified, gel size selected, quantified, pooled, and sequenced on an Illumina HiSeq2500. Sequencing reads were aligned to mm9 using STAR. HOMER findPeaks was used to call peaks/regions in each sample relative to input. HOMER getDifferentialPeaks was used to identify peaks/regions with significantly increased read density induced by LPS. HOMER getDifferentialPeaks was then used to quantify the percentage of LPS-induced peaks/regions that were significantly reduced in read density in hydroxyestrogen-pretreated, LPS stimulated samples. HOMER annotatePeaks.pl was used to make histograms showing read density at LPS-induced peaks/regions across the 3 conditions for both p65 and H3K27ac ChIP-seqs.

### TMRE staining

TMRE (Sigma 87917) was prepared as a 1mM stock in DMSO. TMRE was added directly to cell culture media to macrophages in multiwell tissue culture plates at 37°C (10nM final concentration). 10 min after TMRE addition, cells were treated with EtOH, 4-OHE1, oligomycin (EMD Millipore 495455), or FCCP (Sigma C2920) at indicated concentrations for 20 minutes. Cell were then placed on ice and washed 1x with ice cold PBS, followed by scraping into PBS supplemented with DAPI for flow cytometry analysis.

### RAW matrix-oxGFP macrophages

Cells were transduced with matrix-oxGFP lentivirus encoding oxGFP^13^ with N-terminally fused COX4L mitochondrial matrix targeting sequence and selected with 10μg/mL blasticidin (Thermo Fisher). For experimental use, treatments of RAW matrix-oxGFP macrophages were performed as described, after which cells were put on ice, media aspirated, and cells scraped into PBS supplemented with DAPI for flow cytometry analysis. HSF1 transcriptional inhibitor KRIBB11 (EMD Millipore 385570) was prepared as a 10mM stock in DMSO and used at indicated concentrations.

### mtDNA/gDNA ratio

RAW macrophages in 24-well plates were treated with EtOH or 5μM 4-OHE1 for indicated times, then lysed directly in 100μL total DNA isolation buffer (10?mM Tris-HCl pH 7.5, 50?mM NaCl, 6.25?mM MgCl2, 0.045% NP-40, 0.45% Tween-20). Lysate was moved to PCR tube, supplemented with Proteinase K (NEB), and incubated 1?h at 56°C. Proteinase K was inactivated by incubation at 95°C for 15?minutes, and 5μL of lysate was used in qPCR reactions for mitochondrial DNA (mtDNA, *mt-Cytb* gene locus) and genomic DNA (gDNA, *Actb* gene locus) amplicons using previously described primers^14^ (IDT, see Supplemental Table 6 for sequences).

### MitoTracker Green staining

RAW macrophages or BMDMs were treated with EtOH or estrogens as described. MitoTracker Green (Invitrogen M7514, reconstituted in DMSO) was added directly to cell culture media at a final concentration of 100nM, and mitochondria were labeled for 45 minutes at 37°C. Cells were then placed on ice and washed 1x with ice cold PBS, followed by scraping into PBS supplemented with DAPI for flow cytometry analysis.

### MitoPy1 staining

RAW macrophages and BMDMs in multiwell tissue culture plates were (pre)treated with estrogens and LPS as described. MitoPy1 (Tocris 4428), prepared as a 10mM stock, was added to cell culture media at 1:1000 for direct staining (37°C for 1 hour). Menadione (Sigma M5615) or H2O2 (Fisher H325) were added the last 10-15 minutes of MitoPy1 staining as positive controls. To harvest, cells were placed on ice and washed 1x with ice cold PBS, followed by scraping into PBS supplemented with DAPI for flow cytometry analysis.

### JC-9 staining

RAW macrophages were treated as described, and JC-9 was added directly to media to a final concentration of 500nM. Cells were stained for at 37°C for 30 minutes, with FCCP addition as a positive control to decrease mtMP. To harvest, cells were placed on ice and washed 1x with ice cold PBS, followed by scraping into PBS supplemented with TOPRO3 to exclude dead cells for flow cytometry analysis. Using an LSR Fortessa analyzer, JC-9 emission after excitation with 405nm (violet) laser was collected in two channels: red BV605 channel (595 LP, 610/20 BP filters), and green AmCyan channel (495 LP, 525/50 BP filters). The JC-9 ratio (red aggregate/green monomer) was calculated and plotted. FCCP-driven decrease in this ratio confirmed the dissipation of mtMP.

### roGFP RAW macrophages

RAW macrophages were transduced with lentiviral constructs encoding for N-terminal fusions of roGFP to COX4L targeting sequence (matrix-roGFP), LACTB targeting sequence (inner membrane space- or IMS-roGFP), or nuclear export sequence (cyto-roGFP), and selected with 10ug/mL puromycin (Thermo Fisher). For experimental use, treatments were performed as described, after which cells were put on ice, media aspirated, and cells scraped into PBS supplemented with DAPI. Using an LSR Fortessa analyzer, roGFP emission after excitation with 405nm (violet) and 488nm (blue) lasers was collected using a 505nm longpass:525/50 bandpass filter combination coupled to each respective laser line. H2O2 (Fisher H325) and DTT (Fisher BP172-5) treatment were used to confirm ability to detect roGFP oxidation and reduction.

### Menadione toxicity/mitochondrial superoxide resistance assay

RAW macrophages or BMDMs were treated overnight (18-24 hours) with EtOH vehicle control or 5μM estrogens. The next day, cells were treated with DMSO or 50μM menadione for a short time period (2-6 hours) before harvest. Cells were moved to ice, media aspirated, and cells scraped into PBS supplemented with DAPI. Cell viability was assessed and quantified as a percentage of total events collected for each sample. Viable cells were defined as “live scatter” gate positive, DAPI negative events, with these gates defined in control EtOH/DMSO-treated samples.

### Seahorse respirometry

For mitochondrial stress test, RAW macrophages and BMDMs were plated in XF24 microplates (50K per well) in a small volume of media (100μL) to promote attachment. After attachment (5-6 hours), media volume was brought to 250μL, and cells were treated overnight (18-24 hours) with EtOH, 5μM 4-OHE1, 100ng/mL LPS, or both. The next day, fresh XF Assay media (supplemented with 1mM sodium pyruvate and 25mM glucose) was prepared and brought to pH 7.4. Cells were washed 1x with XF Assay media, 500μL of XF assay media was added to each well, and plate incubated at 37°C (no CO2) for 45-60 minutes. During this time, XF24 sensor cartridge (in calibrant overnight at 37°C) was loaded with Oligomycin, FCCP, and Antimycin A (Sigma A8674), and Rotenone (Cayman 13995), and loaded into Agilent Seahorse XF24 analyzer for calibration. After calibration, cells were loaded for mitochondrial stress test with sequential injections of mitochondrial inhibitors (all at 1.5μM final concentration). All data presented is raw data/unnormalized, as protein quantification after stress test revealed no significant differences between the various treatments. For measurement of 4-OHE1’s acute effects on basal oxygen consumption, cells were plated and moved to fresh XF assay media as described, and treated with EtOH or 4-OHE1 immediately prior to loading into Seahorse analyzer. Data were analyzed using Wave v2.4 and extracted for plotting in GraphPad Prism 8.

### Statistical analysis

All statistical analysis for qPCR and flow cytometry experiments was performed using GraphPad Prism 8 software. Metabolomics statistical analysis was performed using R scripts available upon request.

### Illustrations

Images were created using BioRender.

### Data Availability

All data sets generated and analyzed in the current study are available from the corresponding authors upon request. RNA-seq and ChIP-seq data will be deposited at the Gene Expression Omnibus (GEO), and accession number provided prior to publication.

